# Structural Basis for CAL1-Mediated Centromere Maintenance

**DOI:** 10.1101/723213

**Authors:** Bethan Medina-Pritchard, Vasiliki Lazou, Juan Zou, Olwyn Byron, Juri Rappsilber, Patrick Heun, A. Arockia Jeyaprakash

## Abstract

Centromeres are microtubule attachment sites on chromosomes defined by the enrichment of CENP-A-containing nucleosomes. To preserve centromere identity, CENP-A must be escorted to centromeres by a CENP-A-specific chaperone for deposition. Despite this essential requirement, many eukaryotes differ in the composition of players involved in centromere maintenance highlighting the plasticity of this process. In humans, CENP-A recognition and centromere targeting is achieved by HJURP and the Mis18 complex, respectively. Here, using crystal structures, we show how *Drosophila* CAL1, an evolutionarily distinct CENP-A chaperone, targets CENP-A to the centromere receptor CENP-C without the requirement of the Mis18 complex: while the N-terminal CAL1 fragment (CAL1_1-160_) wraps around CENP-A/H4 through multiple physical contacts, the C-terminal CAL1 fragment (CAL1_893-914_) directly binds CENP-C cupin dimer. Our work shows CAL1, though divergent at the primary structure, employs evolutionarily conserved and adaptive structural principles to recognise CENP-A/H4 and CENP-C providing insights into the minimalistic principles underlying centromere maintenance.

## Introduction

Centromeres are specialised chromosomal regions that act as a platform for the assembly of kinetochores, the microtubule anchoring sites essential for chromosome segregation during mitosis and meiosis (Musacchio and Desai, 2017). Unlike budding yeast where DNA sequence is sufficient to define centromere identity, in most eukaryotes centromeres are defined by the enrichment of unique nucleosomes containing the histone H3 variant CENP-A (Sekulic and Black, 2012, Zasadzinska and Foltz, 2017). As a consequence, maintenance of CENP-A containing nucleosomes is essential for preserving centromere identity through generations of cell cycles. This is achieved through an epigenetic mechanism that relies on CENP-A as an epigenetic mark (Zasadzinska and Foltz, 2017, Westhorpe and Straight, 2014, McKinley and Cheeseman, 2016, Musacchio and Desai, 2017).

Unlike canonical chromatin maintenance where histone H3 is deposited concomitantly with DNA replication, the centromeric chromatin maintenance is decoupled from DNA replication. As a result, CENP-A levels in the sister chromatids are reduced by half during replication (Hemmerich et al., 2008, Jansen et al., 2007, Mellone et al., 2011, Dunleavy et al., 2009, Lidsky et al., 2013). To ensure stable centromere maintenance, CENP-A nucleosome must be brought back to their original levels through active CENP-A deposition. The timing of CENP-A deposition varies among species, however, the underlying mechanisms appear to share significant similarity (Zasadzinska and Foltz, 2017). A central player in this process is the CENP-A-specific chaperone HJURP in human and its homologue Scm3 in fungi (Kato et al., 2007, Foltz et al., 2009, Dunleavy et al., 2011, Pidoux et al., 2009, Sanchez-Pulido et al., 2009). Both HJURP and Scm3 can bind the CENP-A–Histone H4 (CENP-A/H4) heterodimer in its pre-nucleosomal form and these complexes are then targeted to centromeres by the Mis18 complex (Hayashi et al., 2014, Fujita et al., 2007, Nardi et al., 2016, Stellfox et al., 2016, McKinley and Cheeseman, 2014, Moree et al., 2011, Dambacher et al., 2012, French et al., 2017, Hori et al., 2017). While the human Mis18 complex is composed of Mis18a, Mis18b and Mis18BP1, the fission yeast Mis18 complex consists of Mis18, Mis16, Eic1 and Eic2, where Eic1 and Eic2 are proposed to be a functional equivalents of human Mis18BP1 (Subramanian et al., 2014, Hayashi et al., 2014, Fujita et al., 2007). The timing of Mis18 complex assembly, its centromere targeting and subsequent CENP-A deposition is suggested to be tightly controlled by the kinase activities of CDK and Plk1 (McKinley and Cheeseman, 2014, Silva et al., 2012, French and Straight, 2019, Stankovic et al., 2017). While we know the identity of key players involved in centromere maintenance, molecular and mechanistic understanding of their intermolecular cooperation are just emerging (Spiller et al., 2017, Pan et al., 2017, Nardi et al., 2016, Stellfox et al., 2016).

Strikingly, *Drosophila* species have regional centromeres defined by the presence of CENP-A (also called CID) but lack clear homologues of HJURP and the subunits of the Mis18 complex. Instead, fly-specific CAL1 appears to combine both the roles of HJURP and the Mis18 complex: pre-nucleosomal CENP-A recognition and its targeting to the centromere for deposition, respectively (Phansalkar et al., 2012). Targeting CAL1 to non-centromeric DNA in *Drosophila* cells recruited CENP-A and established centromeres capable of assembling kinetochore proteins and microtubule attachments (Chen et al., 2014). These observations and the ability of CAL1 to bind CENP-A/H4 and CENP-C with its N- and C-terminal regions, respectively, collectively established CAL1 as a ‘self-sufficient’ CENP-A-specific assembly factor in *Drosophila* (Chen et al., 2014, Schittenhelm et al., 2010). However, structural level mechanistic understanding of how CAL1 binds CENP-A/H4 and CENP-C to facilitate the establishment and maintenance of centromeres is yet to be determined. The simplistic nature of the centromere maintenance pathway in *Drosophila* makes it a unique model system to understand the fundamentally conserved structural principles underlying centromere maintenance.

In this study, we present the structural basis for the recognition of CENP-A/H4 and CENP-C by CAL1. Our analysis reveals that although CAL1 does not share noticeable sequence similarity with its human or fission yeast counterpart, it recognises CENP-A/H4 using conserved structural principles. We also provide the structural framework of interactions responsible for CENP-C recognition by CAL1. The structural analysis, together with validation of structure-guided mutants *in vitro* and in cells, provides the structural basis for the mechanism by which CAL1 solely recognises and targets CENP-A to centromeres to maintain centromere identity in flies.

## Results

### The N-terminus of CAL1 forms a heterotrimer with the histone fold domain of CENP-A and H4

Secondary structure prediction analysis indicated that CAL1 is likely to be a predominantly unstructured protein, although it includes an N-terminal domain spanning amino acid (aa) residues 1-200 predicted to fold into a helices (Figure S1A and B). With the aim of structurally characterising the intermolecular interactions responsible for CAL1 binding to CENP-A/H4, we reconstituted a protein complex containing the N-terminal 160 aa residues of CAL1 (CAL1_1-160_), a putative histone fold domain of CENP-A (CENP-A_101-225_) and H4 (His-CAL1_1-160_–CENP-A_101-225_–H4) (Figure 1A) using recombinant proteins as previously reported (Chen et al., 2014). Limited proteolysis experiments performed on CAL1_1-160_–CENP-A_101-225_–H4 complex using different proteases suggested that a CENP-A fragment containing aa residues 144-255 (CENP-A_144-255_) is sufficient to interact with CAL1 and H4 (results not shown). Subsequently, using CAL1_1-160_, CENP-A_144-255_ and H4 we reconstituted a truncated protein complex (His-CAL1_1-160_–CENP-A_144-225_–H4). The molecular weights measured for His-CAL1_1-160_–CENP-A_101-225_– H4 and His-CAL1_1-160_– CENP-A_144-225_–H4 using <u>S</u>ize <u>E</u>xclusion <u>C</u>hromatography combined <u>M</u>ulti <u>A</u>ngle <u>L</u>ight <u>S</u>cattering (SEC-MALS) are 47.0 kDa and 43.4 kDa, respectively (Figure S1C). These values match with calculated molecular weights for a 1:1:1 hetero trimeric assembly for both complexes (46.7 kDa and 41.7 kDa for His-CAL1_1-160_–CENP-A_101-225_–H4 and His-CAL1_1-160_– CENP-A_144-225_–H4, respectively) and in agreement with our previous report (Roure et al., 2019). This observation is in striking agreement with the subunit stoichiometry of the human pre-nucleosomal CENP-A/H4 in complex with HJURP (Hu et al., 2011).

**Figure 1 –.**
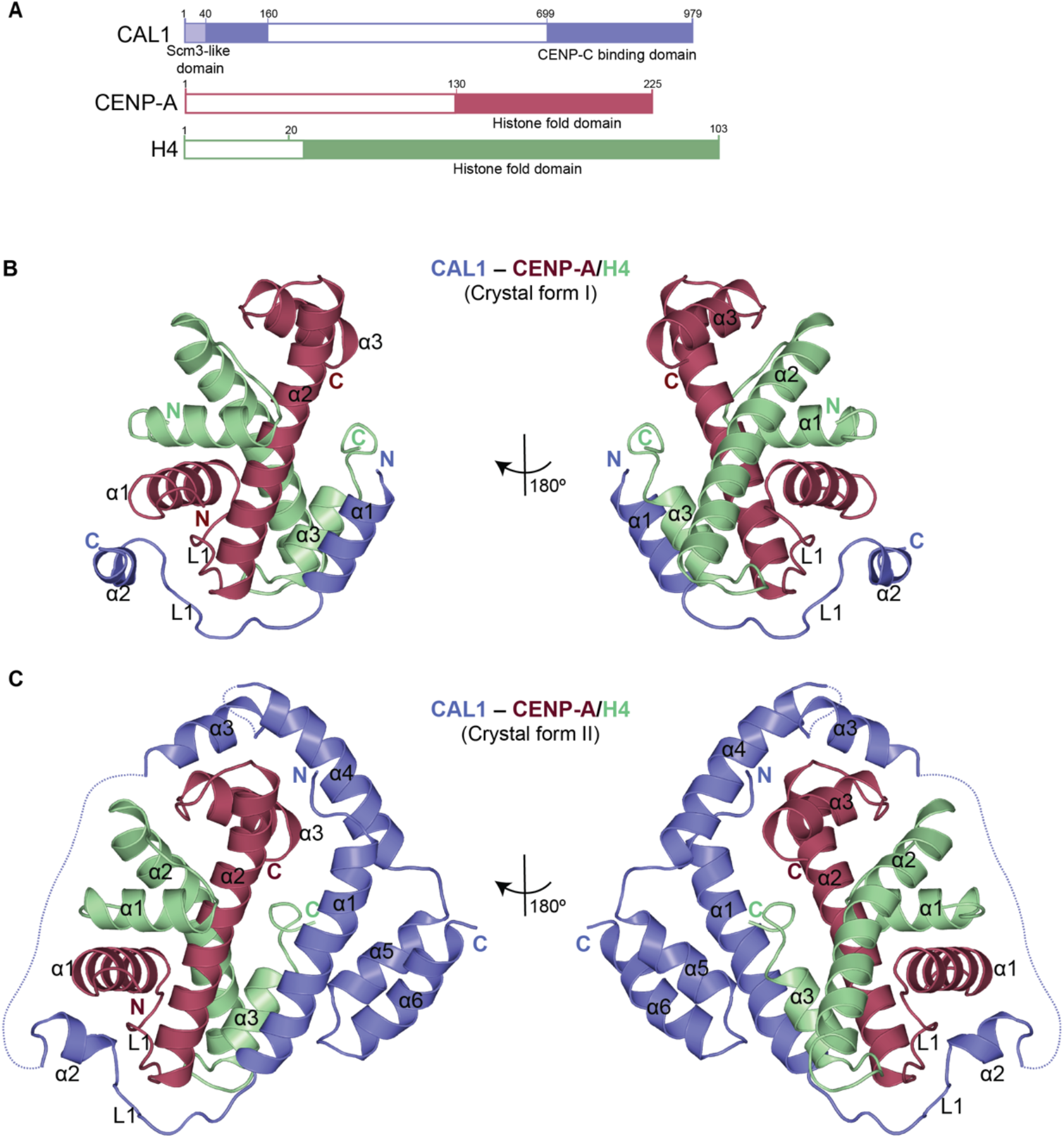
N-terminal 160 amino acids of CAL1 wrap around CENP-A/H4 heterodimer to form a hetero-trimeric assembly. A - Schematic representation of structural features of CAL1, CENP-A and H4. Filled boxes represent folded domains. B - Overall structure of His-CAL1_1-160_–CENP-A_101-225_–H4 (crystal form I). CAL1 is shown in blue, CENP-A in maroon and H4 in green. C - Overall structure of His-CAL1_1-160_–CENP-A_144-225_–H4 (crystal form II). CAL1 is shown in blue, CENP-A in maroon and H4 in green. See also Figures S1-S4

### Structure determination of CAL1_1-160_–CENP-A/H4 complex

Extensive crystallisation trials with CAL1_1-160_–CENP-A_101-225_–H4 and CAL1_1-160_–CENP-A_144-225_–H4 yielded two different crystal forms, form I and form II, that diffracted X-rays to about 3.5 Å (form I) and 4.5 Å (form II) (Table S1). Molecular replacement was performed for the dataset collected from form I using the coordinates of *Drosophila melanogaster* (*dm*) H3/H4 heterodimer (deduced from the structure of *dm* nucleosome core particle, pdb 2PYO) (Clapier et al., 2008). Molecular replacement solution yielded initial phases sufficient for subsequent rounds of model building and refinement (Figure S2A). The final model included residues 18 to 46 of CAL1, 147 to 220 of CENP-A and 28 to 98 of H4 and was refined to an R factor 25.5% and R_free_ factor 28.4% (Figure 1B, Table S1). Although we used a CAL1 fragment spanning residues 1-160 in the crystallisation experiment, the calculated electron density map accounted only for CAL1 residues 14-48. Considering these crystals took more than a year to form, we concluded that CAL1 was proteolytically cleaved, which may have facilitated the crystallisation of a truncated complex.

The refined model obtained using crystal form I was used as a template in molecular replacement to determine the structure of crystal form II (Figure 1C and S2B). The difference electron density map calculated using the molecular replacement solution revealed unambiguous density for most main chain atoms of CAL1_1-160_. While side chain electron densities are well defined for CAL1 residues 7 to 47, only the main chain could be modelled for rest of the CAL1. Considering the modest resolution of the structure, intermolecular interactions stabilising the CAL1–CENP-A/H4 complex were further analysed using chemical <u>C</u>ross-<u>L</u>inking <u>M</u>ass <u>S</u>pectrometry (CLMS). Purified recombinant CAL1_1-160_-CENP-A_101-225_-H4 complex was crosslinked using EDC, a zero-length crosslinker that covalently links carboxylate groups of Asp or Glu residues with primary amines of Lys and N-terminus, or hydroxyl group of Ser, Thr and Tyr. The crosslinked peptides obtained from the trypsin digestion of the crosslinked sample were analysed by mass spectrometry to identify intra-and intermolecular contacts (Figure S3). Notably, the data revealed several intramolecular crosslinks between the N- and C-terminal regions of CAL1_1-160_, suggesting a direct interaction between these regions (Figure S3). This information was particularly helpful in tracing the backbone atoms of residues beyond CAL1 residue 47 within the electron density map.

### Overall structure of CENP-A/H4 assembly

The structures obtained from two different crystal forms together provide key insights into the overall architecture of the assembly (Figure 1B and C). CENP-A residues 147 to 220 form the histone fold domain with characteristic α1, α2 and α3 helices formed by residues 150-164, 174-201 and 207-220, respectively. The corresponding α1, α2 and α3 residues of histone H4 are 30-44, 51-78 and 82-94, respectively. Structural superposition analysis showed that CENP-A/H4 heterodimer aligns well with H3/H4 heterodimer with a root mean square deviation (rmsd) of 1.2 Å (Figure S4A). This suggests that both H3 and CENP-A use an identical mode of H4 binding. However, CENP-A α1, H4 α3 and C-terminal tail show conformational variations in the CAL1-bound CENP-A/H4 complex, likely due to CAL1 binding (Figure S4B). Particularly, in the H3/H4 structure, the C-terminal tail of H4 folds back and makes contacts with the H3 α3 residues close to H3 loop L1, resembling CAL1 interaction at the equivalent region of CENP-A in the CAL1-bound CENP-A/H4 structure. The H4 C-terminal tail possibly swings away from this site upon CAL1 binding. Overall structure of *dm* CENP-A/H4 is very similar to human CENP-A/H4 as these structures superpose well with a rmsd of 1.1 Å (Figure S4C). However, noticeable conformational variation is seen in loop L1, possibly to accommodate the amino acid variations between HJURP and CAL1 (Figure S4B and S4C).

### CAL1 binds CENP-A/H4 heterodimer through multiple physical contacts

CAL1_1-160_ is almost entirely made of a helices that make multiple contacts with CENP-A/H4 heterodimer by wrapping around it (Figure 1C and 2A). Most CENP-A contacts are made by CAL1 helices α1 and α2 and loop L1, which interact with the CENP-A helices α2, α1 and loop L1 respectively involving a total interface area of about 940 Å^2^. Particularly, while the N-terminal half of CAL1 α1 packs against CENP-A α2 involving electrostatic (CAL1 R18 with CENP-AQ90) and hydrophobic (involving CAL1 L11 and M14) interactions, the C-terminal half, mainly aa residues W22 and F29 are sandwiched between CENP-A α2 and H4 α3 (Figure 2A). CAL1 L1 crosses over CENP-A L1 to facilitate CAL1 α2 interaction with CENP-A α3. In addition, CAL1 α4 contacts both CENP-A α2 and α3 involving an interface area of about 80 Å^2^. These CAL1–CENP-A interactions appear to be further stabilised by CAL1 α5 and α6 which together with CAL1 α1 make an intramolecular helical bundle resembling a latch that restrains the position of α1 helix (Figure 1C).

**Figure 2 –.**
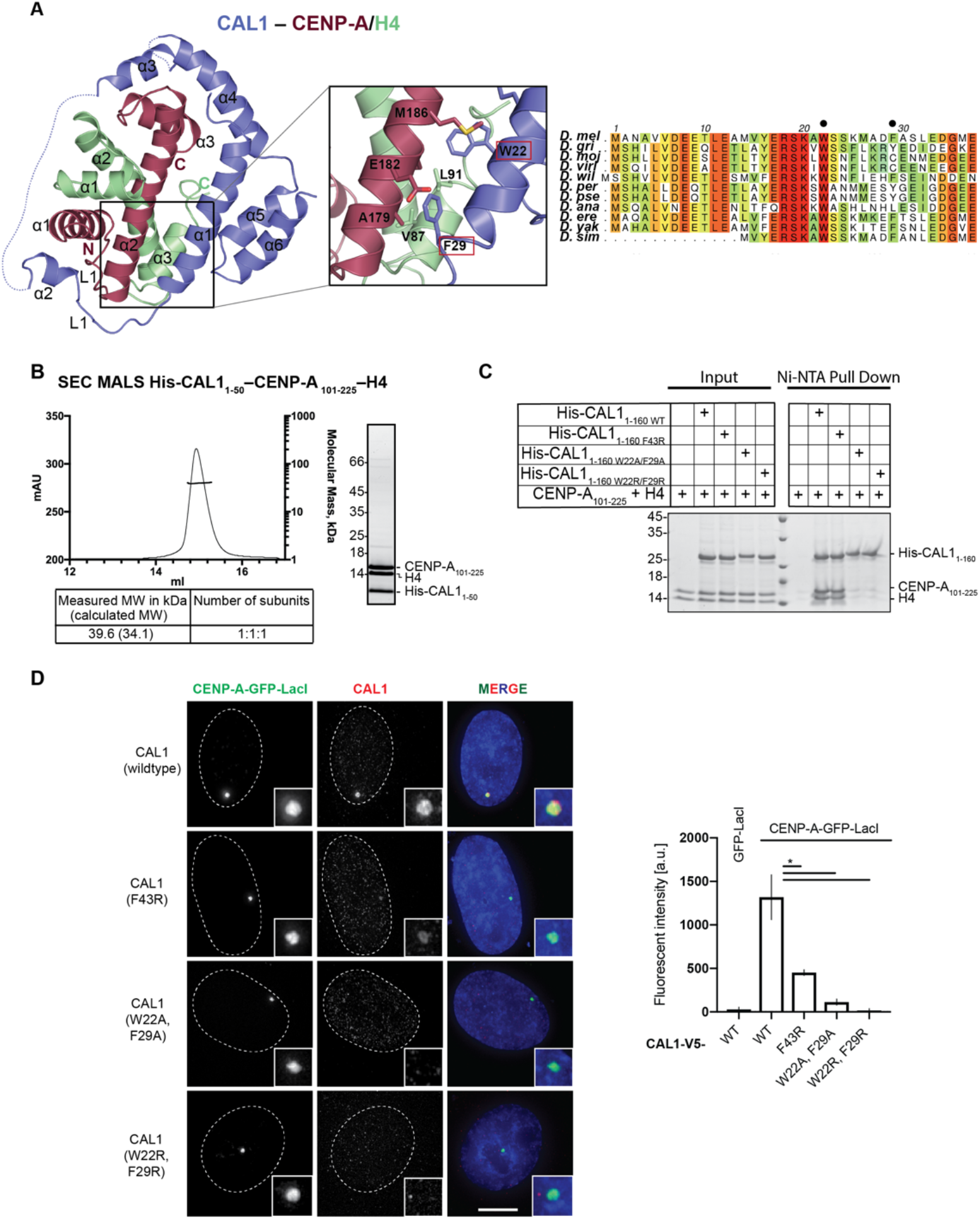
Hydrophobic interactions between CAL1 α1 and CENP-A α1 are critical for CENP-A/H4 binding. A - Crystal structure of His-CAL1_1-160_–CENP-A_144-225_–H4 highlighting residues involved in electrostatic and hydrophobic interactions. CAL1 is shown in blue, CENP-A in maroon and H4 in green. Alignment performed with MUSCLE (Madeira et al., 2019) showing conservations of CAL1 homologues in different fly species. Numbers correspond to *D. mel*. *D. melanogaster* (*D. mel*), *D. grimshawi* (*D. gri*), *D. mojavensis* (*D. moj*), *D. virilis* (*D. viri*), *D. willistoni* (*D. wil*), *D. persimilis* (*D. per*), *D. pseudoobscura pseudoobscura* (*D. pse*), *D. ananassae* (*D. ana*), *D. erecta* (*D. ere*), *D. yakuba* (*D. yak*) and *D. simulans* (*D. sim*). B - SEC-MALS of His-CAL1_1-50_–CEN P-A_101-225_–H4. Absorption at 280 nm (mAU, left y-axis) and molecular mass (kDa, right y-axis) are plotted against elution volume (ml, x-axis). Measured molecular weight (MW) and the calculated subunit stoichiometry based on the predicted MW of different subunit compositions. Samples run on a Superdex 200 increase in 50 mM HEPES pH8.0, 2 M NaCl and 1 mM TCEP. C - Ni-NTA pull-down of His-CAL1_1-160 WT_ and indicated mutants with CENP-A_101-225_–H4. SDS-PAGE shows input and protein bound to beads. D - Representative fluorescence images and quantification of *in vivo* tethering assays. U2OS cells containing a LacO array where co-transfected with CENP-A-GFP-LacI with CAL1_WT_-V5 and also with CAL1-V5 carrying point mutations. Graph shows average of 2 experiments, error bars on graph represent standard deviation. n ≥ 60, scale bar is 10 μm (* P<0.05, **P<0.01).

### Hydrophobic interactions involving CAL1 W22 and F29 are critical for CENP-A/H4 binding

Considering the extent of contacts made by the N-terminal 50 aa residues of CAL1, we checked whether CAL1_1-50_ is sufficient to interact with CENP-A/H4. Using bacterially expressed His-CAL1_1-50_, H4 and CENP-A_101-225_, we confirmed complex formation (Figure 2B). Further characterisation using SEC-MALS showed that CAL1_1-50_–CENP-A_101-225_–H4 is a 1:1:1 complex with a measured molecular weight of 39.6 kDa (calculated molecular weight 34.1 kDa) (Figure 2B).

Within CAL1 the conserved residues in α1: W22 and F29, and in α2: F43 are completely buried in the complex, so we hypothesised that these interactions are crucial for CENP-A/H4 binding (Figure 2A). To test this, we produced recombinant His-CAL1_1-160_ carrying either F43R, W22A/F29A or W22R/F29R mutations and tested their ability to interact with CENP-A/H4 complex in a nickel-NTA pull down assay. His-CAL1_1-160_ was mixed with molar excess of CENP-A/H4 complex, then incubated with nickel-NTA, to capture the His-CAL1_1-160_ and any proteins bound to it. Beads were then washed to remove any unbound proteins. Analysis by SDS-PAGE revealed that while the F43R mutation had no effect on CAL1_1-160_ binding, the W22 and F29 mutations reduced the ability of CAL1_1-160_ to capture CENP-A/H4 compared with the WT protein (Figure 2C).

To validate the requirement of these interactions *in vivo*, we expressed CENP-A-GFP-LacI in U2OS cells containing a synthetic array with a LacO sequence integrated in a chromosome arm (Janicki et al., 2004), and analysed its ability to recruit CAL1-V5 (Roure et al., 2019). When CENP-A-GFP-LacI was tethered to the LacO site, CAL1_WT_ was able to recruit CENP-A to the LacO site (Figure 2D). CAL1_F43R_ caused a 3-fold reduction in the levels co-localising with CENP-A. However, when CAL1_W22/F29_ mutants were used for *in vivo* tethering, they showed very little (Ala mutant) or no localisation (Arg mutant) to the LacO with CENP-A-GFP-LacI, suggesting that these residues are crucial for CENP-A deposition in the cell.

### CAL1 uses conserved and adaptive interactions to recognise *Drosophila* CENP-A/H4

Structural superposition of CAL1–CENP-A/H4 onto its respective human and *Kluyveromyces lactis* structures, HJURP–CENP-A/H4 (PDB: 3R45) (Hu et al., 2011) and Scm3–CENP-A/H4 (PDB: 2YFV) (Cho and Harrison, 2011) showed that CAL1 employs a broadly similar mode of CENP-A recognition with a few striking differences (Figure 3A). All CENP-A chaperones compared here use their α1 helix to interact with α2 of CENP-A in an anti-parallel fashion, occluding the tetramerisation of CENP-A/H4 heterodimers. However, in CAL1 the upstream segment of α1 swings away from CENP-A as compared with its counterpart in HJURP and Scm3. Structural superposition-based sequence alignments showed a key amino acid variation in *dm* CENP-A at position 86 as compared with human and yeast CENP-A: Ala is replaced with Met, an amino acid with a long side chain, which appears to push CAL1 α1 away from it. This apparent weakening of CAL1 α1 – CENP-A α2 interaction is likely to be compensated by CAL1 α5 and α6 which together restrains the position of α1 helix by forming a helical bundle. Notably, loop L1 of both CAL1 and HJURP interacts with CENP-A L1 through main chain hydrogen bonding interactions. However, the secondary structural element downstream of L1 that interacts with the hydrophobic groove formed by CENP-A α1 and α2 is a three stranded β sheet in HJURP whilst it is an α helix in CAL1. Strikingly, unlike other histone chaperones, CAL1 shields CENP-A α3 through downstream α helical elements (Figure 1 and 3). This intermolecular interaction appears to be critical for CENP-A recognition as a CENP-A chimera where CENP-A α3 was replaced with histone H3 α3 failed to associate with centromeres (Roure et al., 2019).

**Figure 3 –.**
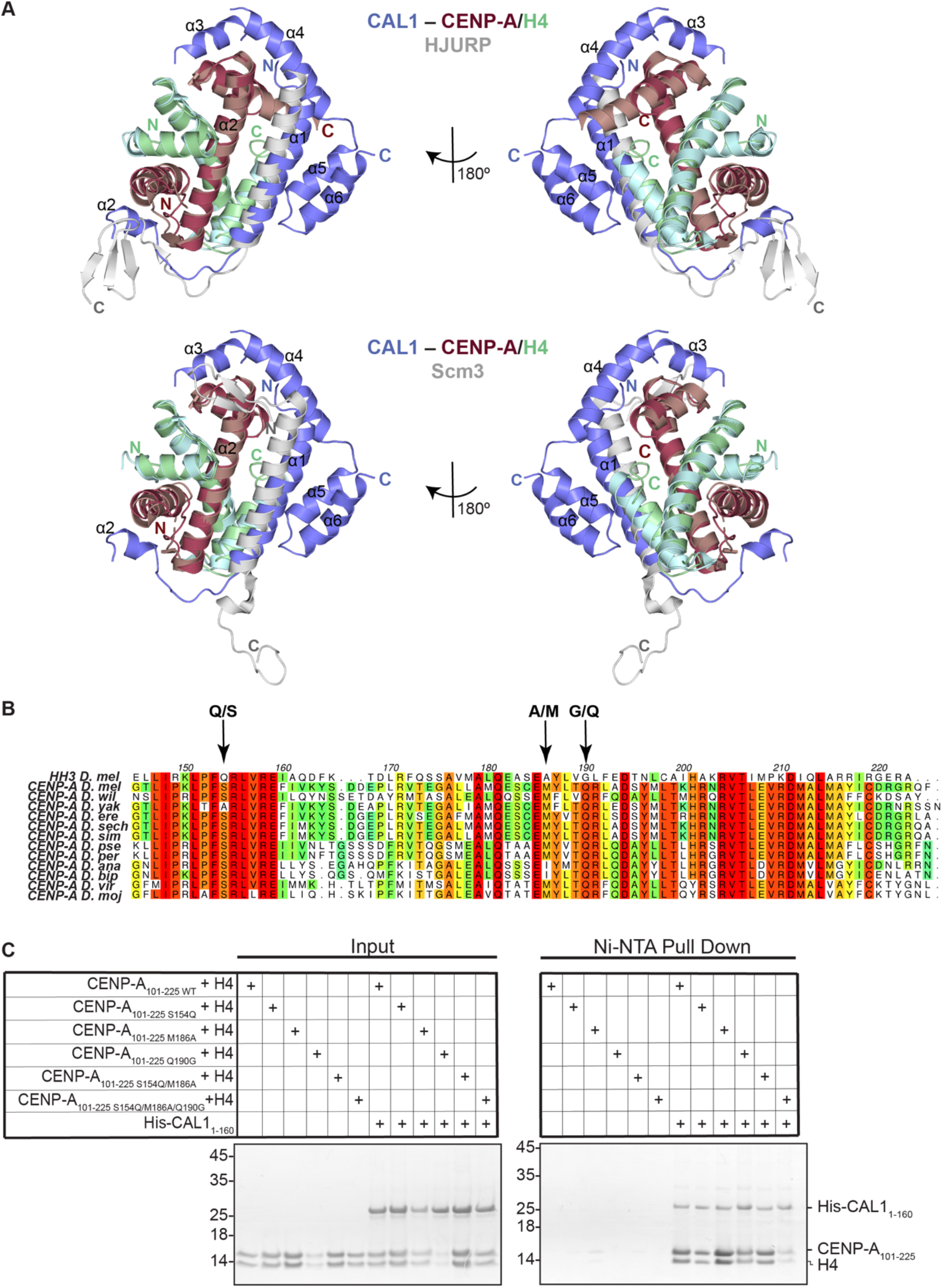
CAL1 uses evolutionarily conserved and adaptive structural interactions to recognise *Drosophila* CENP-A/H4. A - Upper panel shows structure of His-CAL1_1-160_–CENP-A_144-225_–H4 superimposed with structure of human HJURP–CENP-A/H4 (Hu et al., 2011). CAL1 is shown in blue, CENP-A in maroon, H4 in green and HJURP shown in silver. Lower panel shows structure of His-CAL1_1-160_–CENP-A_144-225_–H4 superimposed with structure of yeast Scm3–CENP-A/H4 (Cho and Harrison, 2011). CAL1 is shown in blue, CENP-A in maroon, H4 in green and Scm3 shown in silver. B - Alignment performed with MUSCLE (Madeira et al., 2019) showing conservation of CENP-A homologues in different fly species in comparison to *dm* H3. Numbering corresponds to *D. mel* CENP-A. *D. melanogaster* (*D. mel*), *D. willistoni* (*D. wil*), *D. yakuba* (*D. yak*), *D. erecta* (*D. ere*), *D. sechellia* (*D. sech*), *D. simulans* (*D. sim*), *D. pseudoobscura pseudoobscura* (*D. pse*), *D. persimilis* (*D. per*), *D. ananassae* (*D. ana*), *D. bipectinatal* (*D. bip*), *D. virilis* (*D. viri*), and *D. mojavensis* (*D. moj*). C - Ni-NTA pull-down of His-CAL1_1-160 WT_ with corresponding CENP-A_101-225_–H4 mutants. SDS-PAGE shows input and protein bound to beads.

### CAL1 recognises amino acid variations unique to CENP-A

The histone fold domain of CENP-A and histone H3 share 31% sequence identity. To understand how CAL1 differentiates CENP-A from histone H3 we looked for conserved CENP-A-specific amino acid variations in several *Drosophila* species and compared these variations against *dm* histone H3 (Figure 3B). This analysis together with the structural superposition of CENP-A onto histone H3 revealed several residues unique to CENP-A within the CAL1 binding region potentially responsible for CENP-A specificity: Ser154, Met186 and Gln190. The equivalent residues in histone H3 are Gln, Ala and Gly, respectively. To evaluate if any of these specific amino acid variations is responsible for providing CENP-A specificity, we made several recombinant CENP-A mutants where Ser154, Met186 and Gln190 are mutated to corresponding histone H3 residues Gln (CENP-A_101-225 S154Q_), Ala (CENP-A_101-225 M186A_), and Gly (CENP-A_101-225 Q190G_) and tested the ability of these mutants to interact with His-CAL1_1-160_ in a nickel-NTA pull down assay (Figure 3C). While His-CAL1_1-160_ interacted with CENP-A mutants harbouring single ‘histone H3-like’ mutations as efficiently as it does the WT CENP-A, combining three ‘histone H3-like’ mutations resulted in a noticeable reduction in CAL1 binding (Figure 3C). This suggests that CAL1 achieves CENP-A specificity by recognising multiple CENP-A-specific amino acid variations.

### CAL1 chaperones CENP-A/H4 by shielding protein/DNA interaction surfaces crucial for nucleosome assembly

Histone chaperones are key regulators of nucleosome assembly. This function is achieved by ensuring the correct histone incorporation in a spatio-temporally controlled manner. To understand how CAL1 exerts its CENP-A chaperone function, we performed structural superposition of CAL1–CENP-A/H4 complex onto the crystal structure of nucleosome core particle (PDB: 2PYO) (Clapier et al., 2008). This revealed that CAL1 shields the CENP-A/H4 regions critical for nucleosome assembly at: i) the CENP-A/H4 tetramerisation interface, ii) the H2A/H2B binding region and iii) the DNA-binding region (Figure 4). CENP-A/H4 tetramerisation is thought to be the very first step in the nucleosome assembly pathway, followed by the wrapping of DNA by the CENP-A/H4 heterotetramer and incorporation of H2A/H2B heterodimers (Hammond et al., 2017). Thus, the CAL1 bound form of CENP-A/H4 cannot be incorporated into the nucleosome, inhibiting any unwarranted incorporation of CENP-A.

**Figure 4 –.**
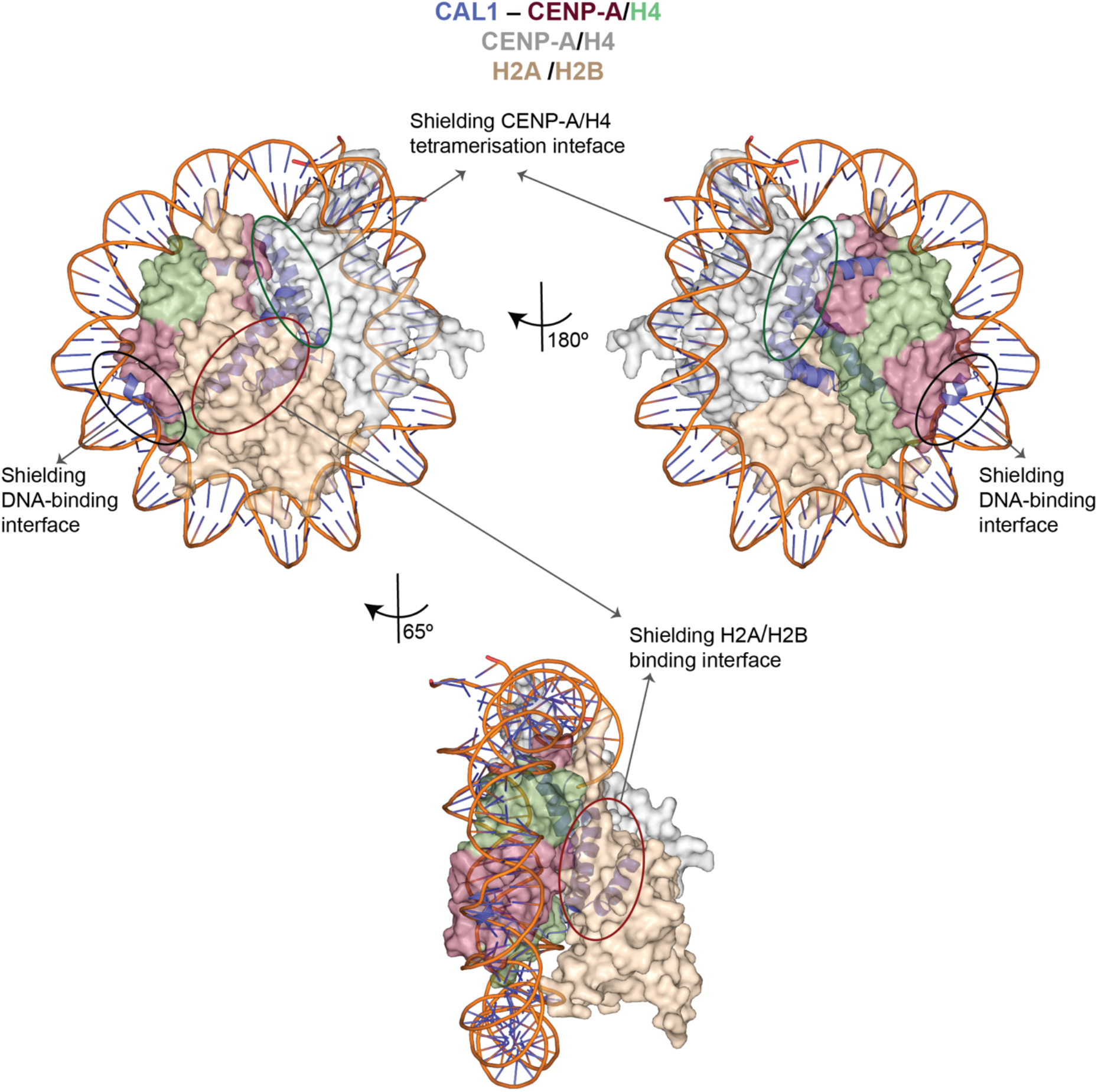
CAL1 chaperones CENP-A/H4 by shielding the CENP-A/H4 tetramerisation, DNA-binding and H2A/H2B-binding interfaces. Structure of His-CAL1_1-160_– CENP-A_144-225_–H4 superimposed with modeled structure of the *dm* CENP-A nucleosome (Clapier et al., 2008). Different orientations highlighting mechanism for specificity to CENP-A/H4 dimer binding by CAL1. CENP-A/H4 and H2A/H2B are shown in surface representation. CAL1 and DNA are shown in cartoon representation. CENP-A and H4 bound to CAL1 are shown in maroon and green while CAL1 in blue.

### CENP-C binds CAL1 via its C-terminal cupin domain

We next aimed to understand the structural basis for the centromere targeting of the CAL1 bound pre-nucleosomal CENP-A/H4 heterodimer. Previous studies have shown that CAL1 and CENP-C can directly interact with each other through their C-terminal regions, CAL1_699-979_ and CENP-C_1009-1411_, respectively (Schittenhelm et al., 2010). However, efforts to purify these recombinant proteins were not successful as they were sensitive to protease contaminants and so were unstable. Based on secondary structure prediction and sequence conservation analysis, we designed shorter constructs of CAL1 and CENP-C, CAL_1841-979_ and CENP-C_1264-1411_. The CENP-C fragment contains an evolutionarily conserved cupin domain. Reconstitution of CAL1–CENP-C complex using individually purified His-SUMO tagged CAL1_841-979_ (His-SUMO-CAL1_841-979_) and His-tagged CENP-C_1264-1411_ (His-CENP-C_1264-1411_) showed clear complex formation (Figure 5A): His-SUMO-CAL_1841-979_ eluted at a volume of 10.38 ml, His-CENP-C_1264-1411_ 10.54 ml, whilst the complex eluted at 9.63 ml.

**Figure 5 –.**
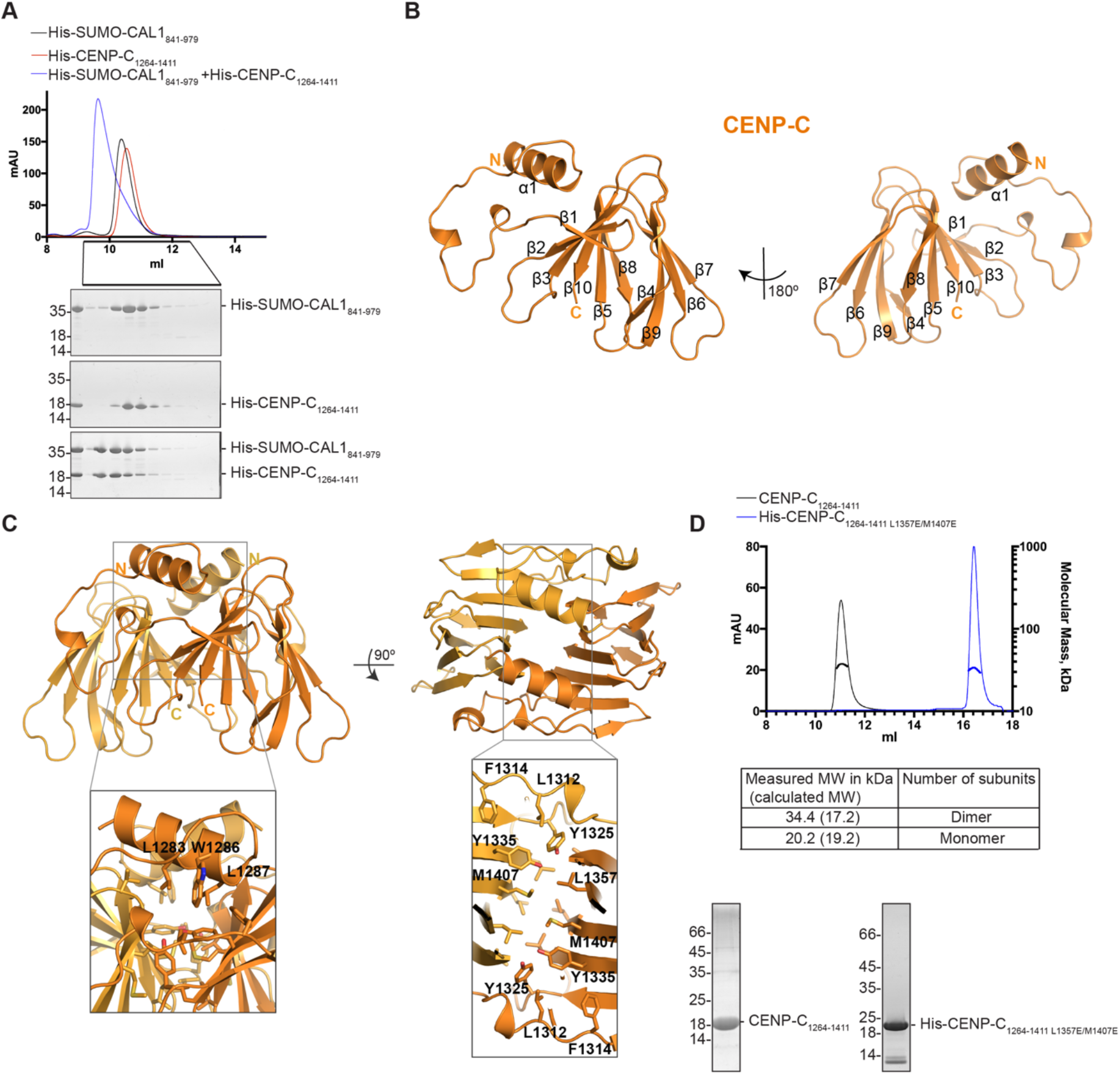
CAL1 binds CENP-C by directly interacting with the evolutionarily conserved Cupin domain. A - SEC profile of His-SUMO-CAL1_841-979_ (black), His-CENP-C_1264-1411_ (red) and His-SUMO-CAL1_841-979_ mixed with molar excess of His-CENP-C_1264-1411_ (blue) and corresponding SDS PAGE analysis of the fractions. Samples run on a Superdex 75 increase 10/300 in 20 mM Tris-HCl pH8.0, 100 mM NaCl and 2 mM DTT. B - Crystal structure of CENP-C cupin domain determined at 1.7Å resolution. C - Overall structure of CENP-C cupin domain dimer. Amino acids residues involved in dimerisation are highlighted in zoomed in panels. D - SEC-MALS analysis of CENP-C_11264-1411_ (black) and His-CENP-C_11264-1411 L1357E/M1407E_ (blue). Absorption at 280 nm (mAU, left y-axis) and molecular mass (kDa, right y-axis) are plotted against elution volume (ml, x-axis). Measured molecular weight (MW) and the calculated subunit stoichiometry based on the predicted MW of different subunit compositions. Samples run on either a Superdex 75 or a Superdex 200 increase 10/300 in 50 mM HEPES pH8.0, 100 mM or 300 mM NaCl and 1 mM TCEP.

### Overall structure of CENP-C cupin domain

A well conserved structural feature of CENP-C among different species is the presence of a C-terminal cupin domain. Previous structural characterisation of the cupin domain of Mif2p, the budding yeast orthologue of human CENP-C, showed that it forms a dimer (Cohen et al., 2008). Although CENP-Cs across species contain a C-terminal cupin domain, these appear to show striking amino acid variations. Pairwise sequence alignments of *Drosophila* CENP-C cupin domain against its budding yeast counterpart showed 11% sequence identity and 18% sequence identity against the human counterpart. Crystallisation trials carried out with CENP-C_1264-1411_ alone and in complex with CAL1 produced diffraction quality crystals which diffracted X-rays to about 1.7 Å and 2.4 Å, respectively (Table S1).

The CENP-C_1264-1411_ structure was determined by molecular replacement using the crystal structure of budding yeast Mif2p cupin domain (PDB: 2VPV) (Cohen et al., 2008). The twofold axis of the CENP-C_1264-1411_ dimer was aligned with the crystallographic two-fold axis. Consequently, just one molecule was present in the asymmetric unit (Figure 5B). As expected, CENP-C_1264-1411_ domain forms a cupin fold almost entirely made of β strands forming a β-barrel with a helix preceding the cupin domain. The β strands assemble into two β sheets: a six-stranded (β1-β2-β3-β10-β5-β8) and a four-stranded (β4-β9-β6-β7) (Figure 5B). The β1 of the six-stranded β sheet is connected to the preceding α1 (spanning aa residues 1276-1288) with a long loop (aa residues 1289-1313) containing two short α helical segments. Dimerisation of CENP-C cupin domain is mediated by a back-to-back arrangement of six stranded β-sheets. In this arrangement the loop connecting the N-terminal α helix (α1) to β1 crosses-over to its dimeric counterpart resulting in a ‘roof’ like positioning of α-helices on top of the β barrels. The surface area buried at the dimerisation interface is 1706 Å^2^ which is about 50% of the total solvent accessible surface area. The interactions stabilising the dimerisation are predominantly hydrophobic involving residues L1283, W1286, L1287, L1312, L1314, Y1325, Y 1335, M1407 and L1357 (Figure 5C). Among these residues, M1407 and L1357 are centrally located and juxtaposed within the hydrophobic core. This led us to hypothesise that these residues may be critical for the assembly of cupin dimer. To test this, we generated a mutant where M1407 and L1357 were mutated to glutamic acids (CENP-C_1264-1411 M1407E/L1357E_) and analysed their oligomeric structure by measuring the molecular weight using SEC-MALS (Figure 5D). While the measured molecular weight of CENP-C_1264-1411_ agreed with the calculated molecular weight of a dimer, the corresponding value for the His-CENP-C_1264-1411_ M1407E/L1357E revealed that it was a monomer (measured molecular weight 20.2kDa and calculated molecular weight 19.2kDa) (Figure 5D).

Structural comparison of *dm* and budding yeast CENP-C cupin domains showed that although these domains share only weak similarity at the amino acid sequence level (21%), the overall fold conferring the β barrel structure is conserved. However, two loop regions (*dm* CENP-C 1324-1333 and 1368-1376) show striking conformational variation as compared with their equivalent regions in budding yeast CENP-C, Mif2p (Figure S5A).

### Structural basis for CAL1 recognition by CENP-C

The structure of CENP-C_1264-1411_ bound to CAL1_841-979_ was determined by molecular replacement using the CENP-C_1264-1411_ structure reported here as a search model. The final model was refined to R and R_free_ factors of 23.0 % and 26.9 %, respectively and included CENP-C residues 1303-1411 and CAL1 residues 890-913 (Figure 6A). This suggests that CAL1 residues preceding and following residues 890 and 913, respectively, are flexible and are not stabilised by CENP-C. While CAL1 residues 890-893 form a β-strand, residues 894-913 form a highly basic α helix (calculated pI of 10.57). CENP-C binds CAL1 using a cradle shaped surface formed by loops L1, L2 and L3 and β-strands β1 and β2. The calculated electrostatic surface properties show that CAL1 binding involves a surface suitable for both electrostatic and hydrophobic interactions (Figure 6A). CAL1 residues 890-893, which form a β-strand interacts with β1 of CENP-C cupin domain running parallel to it and as a consequence extend the β sheet involved in cupin dimerisation. The CAL1 α helix consisting of residues 894-913 makes several hydrophobic (involving L896, I900, W904 and Y908) and electrostatic (R903 and K906) interactions with a complementary hydrophobic (involving residues Y1315, V1317, Y1322 and F1323) and acidic (S1295, E1311 and N1326) region of the cradle shaped CENP-C surface (Figure 6). To evaluate the requirement of these interactions to stabilise CAL1–CENP-C binding, we mutated conserved CENP-C F1324 to Arg (F1324R) and CAL1 I900 to Arg and K907 and Y908 to Ala (I900R/K907A/Y908) and tested the ability of these mutants to bind wild type CAL1 and CENP-C, respectively, in separate SEC experiments (Figure 7A and B). Both His-CENP-C_1264-1411 F1324R_ and His-SUMO-CAL1_841-979 I900R/K907A/Y908A_ failed to interact with His-SUMO-CAL1_841-979_ and His-CENP-C_1264-1411_, respectively and hence eluted at their original elution volumes as compared with the elution volume of the CENP-C– CAL1 complex.

**Figure 6 –.**
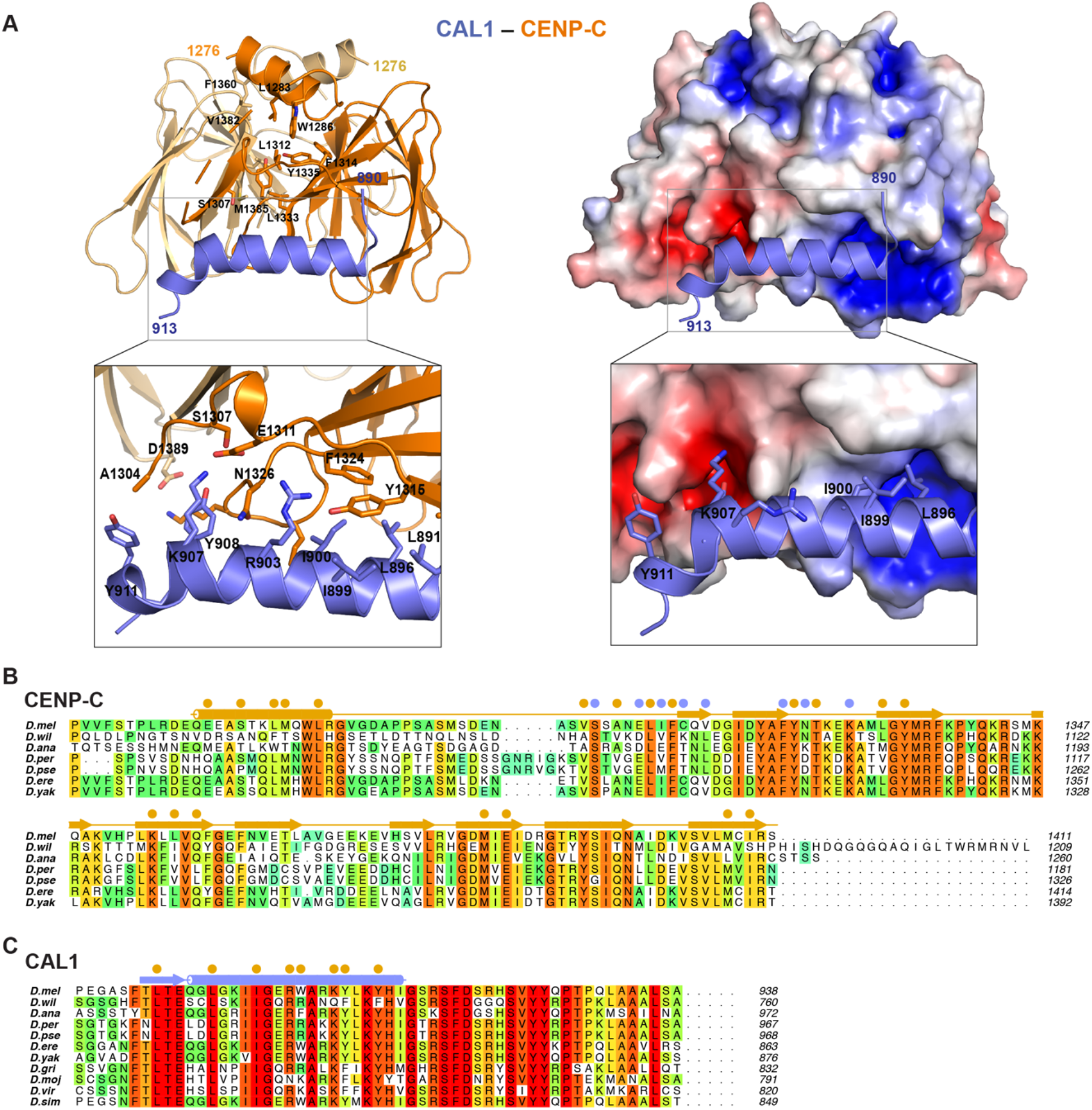
Structural basis for CAL1 recognition by CENP-C cupin domain. A - (left panel) Crystal structure of CAL1 (shown in blue) bound CENP-C cupin domain (shown in orange) determined at 2.4 Å resolution shown in cartoon representation. (right panel) CENP-C cupin domain is shown in surface representation colored based on electrostatic surface potential calculated using APBS. Zoomed in views highlight residues involved in interaction. B - Multiple sequence alignment performed with MUSCLE (Madeira et al., 2019) showing amino acid conservations of CENP-C homologues in different fly species. Residues involved in CENP-C cupin dimerisation and CAL1 binding are highlighted with filled orange and blue circles, respectively. C - Multiple sequence alignment performed with MUSCLE (Madeira et al., 2019) showing conservations of C-terminus of CAL1 across its homologues in different fly species. Orange filled circles highlight the residues involved in CENP-C binding. *D. melanogaster* (*D. mel*), *D. willistoni* (*D. wil*), *D. ananassae* (*D. ana*), *D. persimilis* (*D. per*), *D. pseudoobscura pseudoobscura* (*D. pse*), *D. erecta* (*D. ere*), *D. yakuba* (*D. yak*), *D. grimshawi* (*D. gri*), *D. mojavensis* (*D. moj*), *D. virilis* (*D. viri*), and *D. simulans* (*D. sim*).

We next evaluated the contribution of CENP-C and CAL1 residues identified here as critical for interaction *in vitro* in cultured cells where LacO arrays are integrated in one of the chromosome arms. Tethering GFP-LacI-CENP-C recruited CAL1-V5 to the LacO array. However, the F1324R or the L1357E/M1407E mutation in CENP-C and I900R/K907A/Y908A mutations in CAL1 are both able to inhibit interaction and reduce co-localisation at the tethering site by 4 to 7-fold (Figure 7D).

**Figure 7 –.**
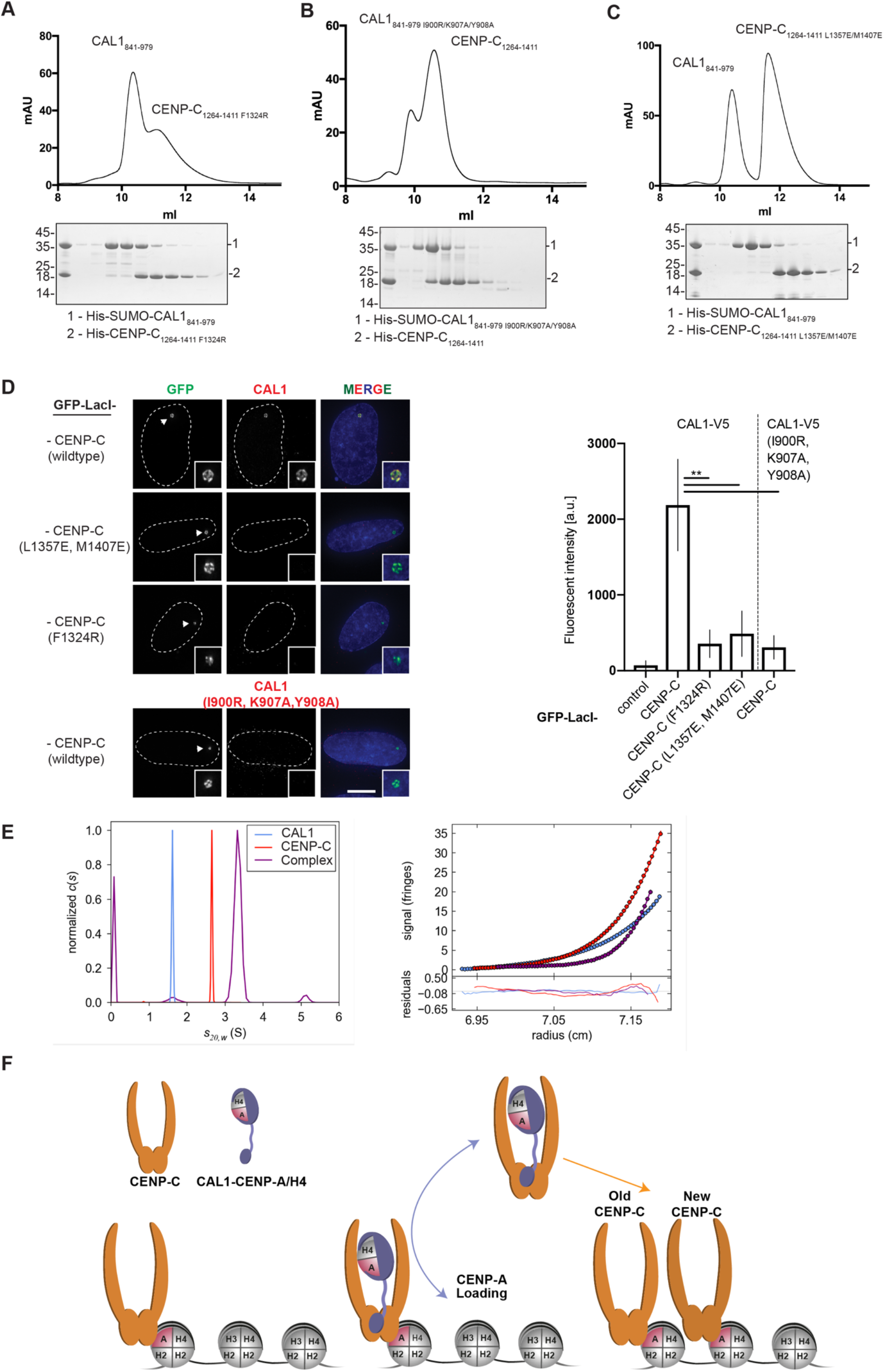
CENP-C cupin dimerisation is critical for CAL1 binding. A - SEC analysis of His-CENP-C_1264-1411 F1324R_ and His-SUMO-CAL1_841-979_ run on a Superdex 75 in 20 mM Tris-HCl pH8.0, 0.1 M NaCl and 2 mM DTT. Corresponding fractions shown by SDS-PAGE with commassie stain underneath. B - SEC analysis of His-CENP-C_11264-1411_ and His-SUMO-CAL1_841-979 I900R/K907A/Y908A_ run on a Superdex 75 in 20 mM Tris pH8.0, 0.1 M NaCl and 2 mM DTT. Corresponding fractions shown by SDS-PAGE underneath. C - SEC analysis of His-CENP-C_11264-1411 L1357E/M1407E_ and His-SUMO-CAL1_841-979_ run on a Superdex 75 in 20 mM Tris pH8.0, 0.1 M NaCl and 2mM DTT. Corresponding fractions shown by SDS-PAGE underneath. D - Representative IF images and quantification of *in vivo* tethering assays. U2OS cells containing a LacO array where transfected with GFP-LacI-CENP-C with CAL1-V5 to assess interaction. CENP-C mutants F1324R, L1357E/M1407E and CAL1 mutant I900R/K907A/Y908A were tested in each construct separately. Graph shows average of 3 experiments, error bars on graph represent standard deviation. n ≥ 136, scale bar is (* P<0.05, **P<0.01) E - (left panel) Normalised sedimentation coefficient distribution (c(s)) for CAL1_841-979_ (CAL1, blue), CENP-C_1264-1411_ (CENP-C, red) and their equimolar mix (Complex, purple) all at 10 mg/ml, demonstrating a significant increase in 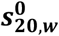 consistent with the formation of a 2:1 complex. (right panel) Typical sedimentation equilibrium data for CAL1_841-979_ (CAL1, blue), CENP-C_1264-1411_ (CENP-C, red) and their equimolar mix (Complex, purple) all at 10 mg/ml, demonstrating a significant increase in mass, consistent with the formation of a 2:1 complex. The data were fit with a single species model yielding masses of 20,253, 29,216 and 49,539 g/mol. The values reported in Figure S6 are based on data acquired for a range of concentrations. See also Figures S5B F - Schematic model of CAL1-mediated loading of CENP-A/H4 and CENP-C at centromeres.

### Dimerisation of CENP-C cupin domain stabilises the CAL1 binding site

Previously we showed that CENP-C dimerisation is required for CAL1 binding in cells (Roure et al., 2019). In the crystal structure presented here, the CAL1 binding site on CENP-C is in close proximity to the cupin dimerisation interface: the loop L1 and β-strands β1, β2 and β3 are all directly involved in stabilising the cupin dimer. This led us to hypothesise that the CAL1 binding site is stabilised in the right conformation by the dimerisation interface and hence disrupting the dimerisation interface might affect CAL1 binding. To test this, we evaluated using SEC the ability of His-CENP-C_1264-1411 M1407E/L1357E_, which we have shown here is not capable of forming a dimer (Figure 5D), to bind CAL1. His-SUMO-CAL1_841-979_ was mixed with 1.2 molar excess of His-CENP-C_1264-1411 M1407E/L1357E_ and subjected to SEC analysis. His-SUMO-CAL1_841-979_ and His-CENP-C_1264-1411 M1407E/L1357E_ did not interact with each other and eluted separately at elution volumes 10.4 ml and 11.6 ml, respectively (Figure 7C). Consistent with these *in vitro* data, GFP-LacI-CENP-C tethered to the LacO site in U2OS cells recruited CAL1-V5 robustly, while the GFP-LacI-CENP-C_M1407E/L1357E_ failed to do so (Figure 7D). These observations together demonstrate that the CENP-C dimerisation-mediated stabilisation of CAL1 binding site is an essential requirement for centromere targeting of CAL1.

### CENP-C cupin dimer binds just one CAL1

Although CAL1 binding by CENP-C involves just a cupin monomer, only one of the two cupin monomers was observed to interact with CAL1 while the equivalent CAL1 binding site of the dimeric counterpart was empty in the crystal structure. We speculate that the other binding site might be sterically hindered by the remaining residues of CAL1 not seen in the crystal structure, thus not allowing a second monomer of CAL1 to bind. This agrees with our previous observation that CAL1_841-979_ and CENP-C_1264-1411_ form a 1:2 complex in solution as estimated using the mass spectrometry derived iBAC peptide ratio and SEC-MALS (Roure et al., 2019). To confirm the subunit stoichiometry of CAL1-CENP-C complex unambiguously, we measured the molecular mass of CAL1_841-979_–CENP-C_1264-1411_ complex using Analytical Ultra Centrifugation (AUC) (Figure 7E). First, the individual components of the complex were characterised by both sedimentation velocity (SV) and sedimentation equilibrium (SE). The data from which (Figure S5B) demonstrate that CAL1_841-979_ is monomeric with a very weak tendency to self-associate, while CENP-C_1264-1411_ is a dimer. Next, samples comprising a complex were analysed. The complex was formed by mixing an equimolar ratio of untagged CAL1_841-979_ and CENP-C_1264-1411_, subjecting it to SEC and using the peak containing the complex for AUC analysis. Both the mass and sedimentation coefficient are consistent with a 2:1 complex, but not with a 2:2 complex. Thus, the AUC together with the crystal structure show that CENP-C cupin dimer binds just one copy CAL1 at any given time.

## Discussion

Understanding the molecular details of how organisms maintain their centromere identity has been of great importance to biologists as loss of centromeres or establishment of new centromeres at non-centromeric locus (neocentromeres) results in genome instability, often leading to cell death. To maintain centromere identity defined by the enrichment of CENP-A containing nucleosome, the CENP-A-specific chaperone (HJURP in humans and Scm3 in yeast) escorts CENP-A until its incorporation into the centromeric chromatin (Pidoux et al., 2009, Foltz et al., 2009, Dunleavy et al., 2009). Correct spatio-temporal regulation of this process is achieved by the Mis18 complex in humans and fission yeast (Fujita et al., 2007, Hayashi et al., 2004, Foltz et al., 2009, Spiller et al., 2017, Pan et al., 2017, McKinley and Cheeseman, 2014). Despite the essential requirement of CENP-A deposition at centromeres, the pathways and the molecular players regulating this process show significant variations across organisms (Zasadzinska and Foltz, 2017). This suggests that these organisms have evolved to employ unique strategies to establish and maintain centromeric chromatin.

*Drosophila* is a remarkable model organism to study centromere inheritance as it lacks direct homologs of either HJURP and Scm3 or the Mis18 complex. Instead it maintains centromere identity using just CAL1. CAL1 does not share obvious sequence similarity with Scm3 or HJURP and does not appear to share common ancestry with these chaperones (Phansalkar et al., 2012, Rosin and Mellone, 2016, Sanchez-Pulido et al., 2009). Our structural analysis presented here shows that although CAL1 appears to have evolved independently of Scm3 and HJURP, it employs evolutionarily conserved structural principles to bind CENP-A. Recognition of CENP-A L1 and α2 by the N-terminal 50 aa residues of CAL1 is remarkably similar to that of Scm3 and HJURP. Despite this structural similarity, CAL1 is also distinctly dissimilar from Scm3 and HJURP as residues downstream of the N-terminal 50 aa wrap around CENP-A–H4 making additional contacts with CENP-A α3 and CAL1 itself. These interactions appear to be crucial for CENP-A deposition as the N-terminal 50 amino acids of CAL1 were not sufficient to recruit CENP-A to centromeres in cells (Chen et al., 2014). Notably, unlike the human CENP-A, the centromere targeting domain of *Drosophila* CENP-A includes α3 as L1 and α2 were not sufficient to target CENP-A (Roure et al., 2019).

The overall mode of CENP-A/H4 recognition by CAL1 appears to be novel as none of the available ‘histone variant’ – chaperone complex crystal structures shows a similar mode of ‘histone variant’ recognition: wrapping around CENP-A/H4 through multiple CENP-A and H4 contacts resulting in the shielding of CENP-A/H4 surfaces involved in CENP-A/H4 tetramerisation, DNA binding and H2A/H2B binding - all critical for nucleosome assembly. This is in agreement with the observation that CAL1 cannot directly interact with the CENP-A nucleosome (Roure et al., 2019) and requires CENP-C to mediate the interaction with the centromeric chromatin.

In humans and fission yeast, the Mis18 complex is responsible for targeting the HJURP bound pre-nucleosomal CENP-A/H4 to the centromere by directly binding CENP-C (reviewed in Westhorpe and Straight, 2014, Zasadzinska and Foltz, 2017, Stellfox et al., 2012) but appears to have been lost during evolution in *Drosophila*. However, CAL1 seems to compensate for this loss by directly associating with CENP-C, which is present in all organisms with monocentric chromosomes (Drinnenberg et al., 2014). While there has been a suggestion that CENP-C cupin domain could be a dimer based on the crystal structure Mif2p cupin domain (Cohen et al., 2008), structural and functional roles of CENP-C cupin domain have remained unclear. Here we show that CAL1 associates with CENP-C by directly interacting with the cupin domain and this interaction is essential for CENP-C mediated recruitment in cells. Our structural analysis shows that the overall structure of *Drosophila* CENP-C cupin domain is similar to that of Mif2p, with striking differences in the mode of dimerisation. It is tempting to suggest that this variation is related to the ability of *Drosophila* CENP-C to bind CAL1 as the CAL1 binding interface of CENP-C is stabilised by dimerisation. Interestingly, although the CENP-C cupin dimer possess two CAL1 binding sites, it appears to accommodate just one CAL1 at a time due to steric hindrance limiting the accessibility of the second CAL1 site. This might have broader implications for the mechanism of CENP-C loading at centromeres (Roure et al., 2019). In the context of the full-length proteins, CAL1 can also oligomerise via its N-terminus (Roure et al., 2019), leading to a scenario where a CENP-C bound CAL1 at the centromere might interact with a second CAL1 bringing another CENP-A/H4 dimer and CENP-C to facilitate CENP-A/H4 tetramer incorporation and the recruitment of CENP-C to the newly formed CENP-A nucleosome (Figure 7F). This is consistent with CENP-C targeting being reliant on CAL1 and CENP-A (Goshima et al., 2007, Erhardt et al., 2008, Schittenhelm et al., 2010, Roure et al., 2019). These observations together with the proposed involvement of HJURP in *de novo* CENP-C recruitment in humans (Tachiwana et al., 2015) suggest that CAL1 is not only a CENP-A loader, but also a CENP-C loader.

In summary, our work demonstrates how *Drosophila* species elegantly compensates for the loss of HJURP or Scm3 and the Mis18 complex through CAL1, which by combining evolutionarily conserved and adaptive structural interactions escorts CENP-A/H4 to the centromere for its subsequent incorporation into the chromatin to maintain centromere identity. Moreover, this is the first study providing the structural basis for how the CENP-A deposition machinery is targeted to centromeres in any organism. Future structural studies on the Mis18 complex and its interaction with HJURP and CENP-C will shed insights into how apparently complex intermolecular interactions achieve the same objective in vertebrates and what are the species-specific functional requirements of this complexity.

## Acknowledgments

We thank the staff of the Edinburgh Protein Production Facility, especially Martin Wear, for their help. Thanks to Atlanta Cook for critical reading of the manuscript. The Wellcome Trust generously supported this work through a Wellcome Trust Career Development Grant (095822) and a Senior Research Fellowship (202811) to AAJ, a Senior Research Fellowship(084229) to JR, a Senior Research Fellowship (103897) to PH, a Centre Core Grant (092076 and 203149) and an instrument grant (091020) to the Wellcome Trust Centre for Cell Biology, a Multi-User Equipment grant 101527/Z/13/Z to the EPPF and a Wellcome-UoE ISSF award toward the procurement of SEC-MALS equipment for the EPPF. PH was further supported by a European Research Council Starting-Consolidator Grant (311674–BioSynCEN).

## Author Contributions

AAJ and PH conceived the project. BM-P, VL, PH and AAJ designed experiments. BM-P,VL, OB and JZ performed experiments. JR provided expertise and feedback. BM-P and AAJ wrote the manuscript with input from all authors.

## STAR*Methods

**KEY RESOURCES TABLE**

| REAGENT or RESOURCE | SOURCE | IDENTIFIER |
| --- | --- | --- |
| Antibodies |  |  |
| Bacterial and Virus Strains |  |  |
| XL1 Blue | Agilent | Cat#200249 |
| Rosetta (*DE3*) | Novagen | Cat#70954 |
| BL21 (*DE3*) pLysS | Novagen | Cat#69451 |
| BL21 Gold (*DE3*) | Agilent | Cat#230132 |
| Biological Samples |  |  |
| Chemicals, Peptides, and Recombinant Proteins |  |  |
| Trypsin | Pierce | Cat#90057 |
| Sulfo-NHS | Thermo Fisher Scientific | Cat#24510 |
| EDC | Thermo Fisher Scientific | Cat#22980 |
| Critical Commercial Assays |  |  |
| Deposited Data |  |  |
| Crystal Structure of CAL1-CENP-A-H4 (form I) | This paper |  |
| Crystal Structure of CAL1-CENP-A-H4 (form II) | This paper |  |
| Crystal Structure of CENP-C | This paper |  |
| Crystal Structure of CAL1-CENP-C | This paper |  |
| Crosslinking Data | This paper |  |
| Experimental Models: Cell Lines |  |  |
| U2OS | (Janicki et al., 2004) |  |
| Experimental Models: Organisms/Strains |  |  |
| Oligonucleotides |  |  |
| All oligonucleotides used in this study are listed in Table S2 | This study | N/A |
| Recombinant DNA |  |  |
| pEC-K-3C-His CAL1<sub>1-160</sub> | This study | N/A |
| pEC-K-3C-His CAL1<sub>1-160</sub> W22A/F29A | This study | N/A |
| pEC-K-3C-His CAL1<sub>1-160</sub> W22R/F29R | This study | N/A |
| pEC-K-3C-His CAL1<sub>1-160</sub> F43R | This study | N/A |
| pEC-K-3C-His CAL1<sub>1-50</sub> | This study | N/A |
| 14S CAL1<sub>841-979</sub> | This study | N/A |
| 14S CAL1<sub>841-979</sub> I900R/K907A/Y908A | This study | N/A |
| pET3a CENP-A<sub>101-225</sub> | Karolin Luger | N/A |
| pET3a CENP-A<sub>101-225</sub> S154Q | This study | N/A |
| pET3a CENP-A<sub>101-225</sub> M186A | This study | N/A |
| pET3a CENP-A<sub>101-225</sub> Q190G | This study | N/A |
| pET3a CENP-A<sub>101-225</sub> S154Q/M186A | This study | N/A |
| pET3a CENP-A<sub>101-225</sub> S154Q/M186A/Q190G | This study | N/A |
| pET3a CENP-A<sub>144-225</sub> | This study | N/A |
| pET22a H4 | Karolin Luger | N/A |
| pEC-K-3C-His CENP-C<sub>1264-1411</sub> | This study | N/A |
| pEC-K-3C-His CENP-C<sub>1264-1411</sub> L1357E/M1407E | This study | N/A |
| pEC-K-3C-His CENP-C<sub>1264-1411</sub> F1324R | This study | N/A |
| pMT CAL1 3Ha Hygro | (Roure et al., 2019) | N/A |
| pMT CAL1 <sub>W22A/F29A</sub> 3Ha Hygro | This study | N/A |
| pMT CAL1 <sub>W22R/F29R</sub> 3Ha Hygro | This study | N/A |
| pMT CAL1 <sub>F43R</sub> 3Ha Hygro | This study | N/A |
| pMT CAL1 <sub>I900R/K907A/Y908A</sub> 3Ha Hygro | This study | N/A |
| pN2 CMV CAL1 V5 | (Roure et al., 2019) | N/A |
| pN2 CMV CAL1 <sub>W22A/F29A</sub> V5 | This study | N/A |
| pN2 CMV CAL1 <sub>W22R/F29R</sub> V5 | This study | N/A |
| pN2 CMV CAL1 <sub>F43R</sub> V5 | This study | N/A |
| pN2 CMV CAL1 <sub>I900R/K907A/Y908A</sub> V5 | This study | N/A |
| pMT_GFP_LacI CENP-C | (Roure et al., 2019) | N/A |
| pMT_GFP_LacI CENP-C <sub>L1357E/M1407E</sub> | This study | N/A |
| pMT_GFP_LacI CENP-C <sub>F1324R</sub> | This study | N/A |
| pN2_CMV_GFP_LacI CENP-C | (Roure et al., 2019) | N/A |
| pN2_CMV_GFP_LacI CENP-C <sub>L1357E/M1407E</sub> | This study | N/A |
| pN2_CMV_GFP_LacI CENP-C <sub>F1324R</sub> | This study | N/A |
| pN2_CMV_CENP-A_GFP_LacI | (Roure et al., 2019) | N/A |
| pMT_3HA CENP-C | (Roure et al., 2019) | N/A |
| pMT_3HA CENP-C <sub>L1357E/M1407E</sub> | This study | N/A |
| pMT_3HA CENP-C <sub>F1324R</sub> | This study | N/A |
| pN2_CMV_3HA CENP-C | (Roure et al., 2019) | N/A |
| pN2_CMV_3HA CENP-C <sub>L1357E/M1407E</sub> | This study | N/A |
| pN2_CMV_3HA CENP-C <sub>F1324R</sub> | This study | N/A |
| **Software and Algorithms** |  |  |
| Clustalw2 v.2.1 | https://doi.org/10.1093/bioinformatics/btm404 | http://www.clustal.org/clustal2/ |
| COOT | (Emsley and Cowtan, 2004) | http://www2.mrc-lmb.cam.ac.uk/personal/pemsley/coot/ |
| PyMOL |  | https://www.pymol.org |
| Prism 7 | GraphPad | https://www.graphpad.com/scientificsoftware/prism/ |
| SEDFIT | Schuck (2000) | http://www.analyticalultracentrifugation.com/default.htm |
| SEDNTERP | Hayes et al. (2012) | http://bitcwiki.sr.unh.edu/index.php/Main_Page |
| SEDPHAT | Vistica et al. (2004) | http://www.analyticalultracentrifugation.com/sedphat/index.htm |
| SOMO | Brookes and Rocco (2018) | http://somo.aucsolutions.com/ |
| Other |  |  |
| cOmplete™-EDTA-free Protease Inhibitor Cocktail | Sigma-Aldrich | Cat#05056489001 |
| HisTrap® HP | GE Healthcare | Cat#17-5248-02 |
| HiTrap® Q HP | GE Healthcare | Cat#17-1154-01 |
| HiTrap® SP HP | GE Healthcare | Cat#17-1152-01 |
| HisPur™ Ni-NTA Resin | Thermo Fisher Scientific | Cat#88222 |
| Superdex 200 increase 10/300 GL | GE Healthcare | Cat#28990944 |
| Superdex 75 increase 10/300 GL | GE Healthcare | Cat#29148721 |
| Phusion Ultra II | Agilent | Cat#600670 |
| T4 DNA Polymerase | NEB | Cat#M0203L |
| Phusion | NEB | Cat#M0530L |
| Bolt™ 4-12% Bis-Tris Plus Gels, 12-well | Invitrogen | Cat#NW04122BOX |
| Bolt™ 4-12% Bis-Tris Plus Gels, 15-well | Invitrogen | Cat#NW04125BOX |
| 20x MES Buffer | Invitrogen | Cat#NP0002 |
| Millex Syringe Filter | Millipore | Cat#SLHVM25NS |
| Crystal Gryphon robot | Art Robbins | N/A |
| JCSG+ | Molecular Dimensions | Cat#MD1-37 |
| Structures Screen 1 + 2 | Molecular Dimensions | Cat#MD1-30 |
| Viscotek SEC-MALS 20 | Malvern Instruments | N/A |
| Viscotek RI Detector | Malvern Instruments | Cat#VE3580 |

## CONTACT FOR REAGENT AND RESOURCE SHARING

Further information and requests for resources and reagents should be directed to and will be fulfilled by the Lead Contact, A. A. Jeyaprakash.

## EXPERIMENTAL MODEL AND SUBJECT DETAILS

### Bacterial Strains

Histones were expressed using BL21 (*DE3*) pLysS, CAL1 using BL21 Gold (*DE3*) and CENP-C using Rosetta *(DE3)* competent *Escherichia coli*.

## METHOD DETAILS

### Plasmids

Codon optimised *Drosophila melanogaster* CAL1 and CENP-C were produced as gBlocks (IDT) with additional end sequences to make them compatible with ligation independent cloning (LIC). CAL1_1-50_, CAL1_1-160_, CAL1_841-979_ and CENP-C_1264-1411_ were produced by either using gBlocks directly or sub-cloning required fragments by Phusion amplification. Inserts and vectors were processed with T4 DNA polymerase according to manual instructions. pET His6 Sumo TEV (14S Addgene plasmid # 48291) was a gift from Scott Gradia. pEC-K-3C-His was a gift from Elena Conti. pET3a CENP-A_101-225_ and pET22b H4 were kind gifts from Karolin Luger.

CAL1_1-160_ mutants (W22A/F29A, W22R/F29R and F43R), CAL1_841-979_ mutants (I900R/K907A/Y908A), CENP-A_101-225_ mutants (S154Q, M186A, Q190G, S154Q/M186A and S154Q/M186A/Q190G) and CENP-C_1264-1411_ mutant (L1357E/M1407E, F1324R) were generated following the site-directed mutagenesis protocol using phusion ultra II (Table S2).

### Protein Production

#### Purification of histones

To purify from inclusion bodies, histones were expressed in *E. coli* BL21 (*DE3*) pLysS cells grown in 2 L of LB media at 37°C until O.D 0.6. Cells were induced with IPTG to a final concentration of 0.2 mM at 37°C for 3-4 hours before harvesting by centrifugation at 4000 x g for 10 min. Pellets were resuspended in 50 mM Tris-HCl pH 8, 100 mM NaCl, 1 mM EDTA, 1 mM benzamidine and 5 mM ßME and snap frozen in liquid nitrogen. After two rounds of freeze thaw cycles the cells were further lysed by sonication and any soluble proteins were separated by centrifugation at 22,000 rpm for 1 h at 4°C. The insoluble pellets were washed twice with 50 mM Tris-HCl pH 8, 100 mM NaCl, 1 mM EDTA, 1 mM benzamidine, 5 mM ßME and 1% triton then twice with 50 mM Tris-HCl pH 8, 100 mM NaCl, 1 mM EDTA, 1 mM benzamidine and 5 mM ßME using a glass homogeniser and pelleting by centrifugation at 22,000 rpm for 10 min at 4°C in between.

Pellets were then left to soak with 500 μl of DMSO for 15 min before resuspending in 20 mM Tris-HCl pH 7.5, 7 M guanidine HCl, 10 mM DTT. After brief sonication, samples were left rotating at RT for 2 h, the unfolded protein was then recovered by centrifugation at RT for 20 min at 22,000 rpm.

Next the histone was dialysed twice for 2 h and once overnight against 500 ml of a buffer containing 10 mM Tris pH 8, 100 mM NaCl, 7 M urea, 1 mM EDTA and 5 mM ßME. Proteins were then centrifuged at 4°C at 22,000 rpm for 30 min, before briefly sonicating then passing through a Millex syringe filter (Millipore). Proteins where then further purified using ion exchange in such a way that samples were passed first through a HiTrap^â^ Q HP column (GE Healthcare) equilibrated with dialysis buffer, then a HiTrap^â^ SP HP column. Protein bound to the HiTrap^â^ SP HP was eluted using a gradient of 10 mM Tris-HCl pH 8, 1 M NaCl, 7 M urea, 1 mM EDTA and 5 mM ßME. After analysis by SDS-PAGE, appropriate fractions were pooled and dialysed twice for 2 h and once overnight against 2 L ddH_2_O and 5 mM ßME. The concentration of the histones was measured by Bradford assay and then proteins were lyophilised for storage.

#### Purification of N-term CAL1 proteins

His-CAL1_1-160_ and His-CAL1_1-50_ were expressed using *E. coli* BL21 (*DE3*) Gold grown in 2XTY media and induced at 18°C using 0.3 mM IPTG before being purified under native and denaturing conditions. Under native conditions pellets were resuspended in a buffer containing 20 mM Tris-HCl (pH 8.0), 100 mM NaCl, 35 mM imidazole and 2 mM ßME and supplemented with cOmplete -EDTA-free (Sigma) before lysing by sonication. Clarified lysates were applied onto a HisTrap^â^ HP column then washed with 60 CV of lysis buffer, 20 CV of 20 mM Tris-HCl (pH 8.0), 1 M NaCl, 35 mM imidazole, 50 mM KCl, 10 mM MgCl_2_, 2 mM ATP and 2 mM ßME then 10 CV lysis buffer. Proteins were then eluted in 20 mM Tris-HCl pH 8.0, 100 mM NaCl, 500 mM imidazole and 2 mM ßME.

For denatured proteins, pellets were suspended in 2 ml/g of wet pellet of 20 mM Tris-HCl pH 8.0, 500 mM NaCl, 25 mM imidazole, 7 M urea and 2 mM ßME, and incubated for 1 h at 4°C with rotation. The suspension was then sonicated to shear DNA before clarifying by centrifugation. Lysate was then incubated with 10 ml of HisPurä Ni-NTA resin (Thermo Fisher Scientific) overnight, before washing with 60 CV of buffer, 20 CV of 20 mM Tris-HCl (pH 8.0), 1 M NaCl, 25 mM imidazole, 7 M urea and 2 mM ßME, 10CV of 500 mM NaCl buffer before eluting with 20 mM Tris-HCl (pH 8.0), 500 mM NaCl, 500 mM imidazole, 7 M guanidine HCl and 2 mM ßME.

#### Protein refolding

To refold histones with and without CAL1, histones were resuspended in 20 mM Tris-HCl pH 7.5, 7 M guanidine HCl and 2 mM ßME and mixed with equimolar amounts of proteins needed. Proteins were then dialysed for 2 h at 4°C against 200 ml of 20 mM Tris-HCl pH 7.5, 7 M guanidine HCl and 2 mM ßME, then 2 L of 10 mM Tris-HCl pH 7.5, 2 M NaCl, 1 mM EDTA and 5 mM ßME was slowly added overnight using a peristaltic pump. If needed, refolded protein was further dialysed against a lower salt concentration solvent; if not, complexes were concentrated and purified by SEC using either a Superdex 200 increase 10/300 GL or Superdex 75 increase 10/300 GL column (GE Healthcare).

#### Purification of CENP-C

His-CENP-C_1264-1411_ was expressed in Rosetta cells using 2XTY and induced at 18°C overnight using 0.3 mM IPTG. Pellets were resuspended in 20 mM Tris-HCl pH 8.0, 100 mM NaCl, 35 mM imidazole and 2 mM ßME and supplemented with 1 mM PMSF and cOmplete-EDTA-free before being lysed by sonication. Clarified lysates were applied to a 5 ml HisTrap^â^ HP column and washed with 80 CV of lysis buffer. Protein was eluted using 20 mM Tris-HCl pH 8.0, 100 mM NaCl, 500 mM imidazole and 2 mM ßME and fractions containing protein were dialysed overnight against 20 mM Tris-HCl pH 8.0, 500 mM NaCl, and 2 mM DTT. Tags were removed by incubation with 3C O/N. Proteins were purified by SEC using either a Superdex 75 10/300 GL or Superdex 75 increase 10/300 GL column (GE Healthcare).

#### Purification of C-term CAL1 protein

His-SUMO-CAL1_841-979_ was expressed in BL21 (*DE3*) Gold in LB and induced at 18°C overnight using 0.3 mM IPTG. Purification was performed as for His-CENP-C_1264-1411_ with an additional ion-exchange step. Dialysed protein was applied to a HiTrap^â^ Q HP column and eluted with a gradient of 10-60% 20 mM Tris-HCl (pH 8.0), 1 M NaCl, and 2 mM DTT over 10 CV. Tags were removed by incubation with TEV overnight followed by a reverse affinity step. Proteins were purified by SEC using either a Superdex 200 increase 10/300 GL or Superdex 75 increase 10/300 GL column (GE Healthcare).

### Crystal Structure Determination and Analysis Crystallisation

Crystallisation trials were preformed using a nanoliter crystal Gryphon robot (Art Robbins) and grown by vapor diffusion methods. For CAL1_1-160_-CENP-A_101-225_-H4, 27 mg/ml of complex in a buffer of 20 mM Tris-HCl pH8.0, 1 M NaCl and 5 mM DTT was used with Structure Screen 1 + 2 (Molecular Dimensions) at 18°C. Crystals were obtained after about a year in G1 – 0.01 M Cobalt (II) chloride hexahydrate, 0.1 M MES pH 6.5 and 1.8 M ammonium sulphate. For CAL1_1-160_-CENP-A_144-225_-H4, protein in 20 mM Tris-HCl pH 8.0, 1 M NaCl and 2 mM DTT was concentrated to 17 mg/ml and used with Structures 1 + 2 at 18°C. Crystals were obtained in C11, 0.2 M lithium sulfate, 0.1 M Tris pH 8.5 and 30% PEG 4000. An optimisation screen was set up in a 24 well format, using half the original concentrations of protein. Tris-HCl pH 8.5 was kept at 0.1 M whilst concentrations of lithium sulfate varied from 0.1-0.3 M and PEG 4000 varied from 24-34%.

Cleaved CENP-C_1264-1411_ was screened at 15 mg/ml in 20 mM Tris-HCl pH 8.0, 500 mM NaCl and 2 mM DTT against several commercial and homemade screens at 18°C. Crystals were obtained in around 13% of all conditions tested. His-CENP-C_1264-1411_-CAL_1841-979_ complex was made in 20 mM Tris-HCl pH 8.0, 100 mM NaCl and 2 mM DTT and used with Structure 1 + 2 and JCSG+ (Molecular Dimensions) at 15 mg/ml at 4°C. Crystals were briefly transferred to a cryoprotectant solution (either oil or the mother liquor supplemented with 40% peg 3350) before directly flash cooled in liquid nitrogen and analysed on beamline i03 and i04-1at the Diamond Light Source (Didcot, UK).

### Data Collection and Crystal Structure Determination

Diffraction data were collected on beamlines i03 (CAL1_1-160_–CENP-A/H4 Form I, CENP-C1_264-1411_; CENP-C_1264-1411_-CAL1_841-979_), i04-1 (CAL1_1-160_–CENP-A/H4 Form II), at the Diamond Light Source (Didcot, UK). Data were processed using the software pipeline available at Diamond Light Source that relies on XDS, CCP4, CCTBX, autoPROC and staraniso (Winter and McAuley, 2011, Kabsch, 2010, Grosse-Kunstleve et al., 2002, Winn et al., 2011, Vonrhein et al., 2011, Tickle et al., 2018). CAL1_1-160_–CENP-A/H4 (Form I and II), CENP-C_1264-1411_, and CENP-C_1264-1411_-CAL1_841-979_ structures were determined by molecular replacement with the program PHASER (McCoy et al., 2007) using the coordinates of *dm* H3/H4 heterodimer deduced from the structure of *dm* nucleosome core particle, PDB: 2PYO (Clapier et al., 2008) and budding yeast Mif2p cupin domain, PDB: 2VPV (Cohen et al., 2008) and *dm* CENP-C_1264-1411_ determined here, respectively. Structures were refined using the PHENIX suite of programs (Adams et al., 2010). CAL1_1-160_–CENP-A/H4 Form II was refined using PHENIX-Rosetta (DiMaio et al., 2013). Model building and structural superpositions were done using COOT (Emsley and Cowtan, 2004). Figures were prepared using PyMOL (http://www.pymol.org). Data collection, phasing and refinement statistics are shown in Table S1.

### Ni-NTA interaction trials

Ni-NTA pull-down assays were performed using His-CAL1_1-160_ WT and mutants mixed with 1.3 times molar excess of CENP-A_101-225_-H4 and made up to 100 μl with 20 mM Tris-HCl pH 8.0, 2 M NaCl, 10% glycerol, 0.5% NP40, 35 mM imidazole and 2 mM ßME. 90 μl was incubated with 60 μl of HisPurä Ni-NTA resin slurry that had been washed with ddH_2_O and buffer for 30 min at 4°C. Beads were then washed four times with 1 ml of buffer, then twice with 1 ml of 20 mM Tris-HCl pH 8.0, 500 mM NaCl, 35 mM imidazole and 2 mM ßME and eluted by boiling in SDS-PAGE loading dye before being separated on a Boltä 4-12% Bis-Tris Plus gel run at 180 V for 1 h in MES buffer.

### SEC-MALS

Size-exclusion chromatography (ÄKTA-Micro™, GE Healthcare) coupled to UV, static light scattering and refractive index detection (Viscotek SEC-MALS 20 and Viscotek RI Detector VE3580; Malvern Instruments) was used to determine the molecular mass of proteins and protein complexes in solution. Injections of 100 μl of 1-5 mg/ml material were used.

For His-CAL1_1-160_-CENP-A_101-225_-H4 (*∂*A_280nm_/*∂*c = 0.67 AU.ml.mg^-1^), His-CAL1_1-160_-CENP-A_144-225_-H4 (*∂*A_280nm_/*∂*c = 0.75 AU.ml.mg^-1^) and His-CAL1_1-50_-CENP-A_101-225_-H4 (*∂*A_280nm_/*∂*c = 0.55 AU.ml.mg^-1^) were run at RT on a Superdex 200 increase 10/300 GL size exclusion column pre-equilibrated in 50 mM HEPES pH 8.0, 2 M NaCl and 1 mM TCEP at 22°C with a flow rate of 0.5 ml/min. His-CENP-C_1264-1411 L1357E/M1407E_ (*∂*A_280nm_/*∂*c = 0.75 AU.ml.mg^-1^) was run on a Superdex 200 increase 10/300 GL size exclusion column pre-equilibrated in 50 mM HEPES pH 8.0, 300 mM NaCl and 1 mM TCEP at 22°C with a flow rate of 0.5 ml/min. CENP-C_1264-1411_ (*∂*A_280nm_/*∂*c = 0.84 AU.ml.mg^-1^) was run at 4°C on a Superdex 75 increase 10/300 GL size exclusion column pre-equilibrated in 50 mM HEPES pH 8.0, 100 mM NaCl and 1 mM TCEP. Light scattering, refractive index (RI) and A280nm were analysed by a homo-polymer model (OmniSEC software, v5.02; Malvern Instruments) using the parameters stated for each protein, *∂*n/*∂*c = 0.185 ml.g^-1^ and buffer RI value of 1.335. The mean standard error in the mass accuracy determined for a range of protein-protein complexes spanning the mass range of 6-600 kDa is ± 1.9%.

### Cross-Linking Mass Spectrometry (CLMS)

Crosslinking was performed on gel filtered complexes dialysied into PBS. 30 μg of zero-length crosslinkers EDC (Thermo Fisher Scientific) and 66 μg sulfo-NHS (Thermo Fisher Scientific) were used to crosslink 10 μg of protein for 1.5 h at RT. The reactions were quenched with 100 mM Tris-HCl before separation on Boltä 4-12% Bis-Tris plus gels (Invitrogen). The bands excised and proteins were reduced with 10 mM DTT for 30 min at room temperature before being alkylated with 55 mM iodoacetamide for 20 min in the dark at room temperature. Proteins were digested with 13 ng/μl trypsin (Pierce) overnight at 37°C. The digested peptides were loaded onto C18-Stage-tips (Rappsilber et al., 2007) for LC-MS/MS analysis. LC-MS/MS analysis was performed using an Orbitrap Fusion Lumos (Thermo Fisher Scientific) with a “high/high” acquisition strategy. The peptide separation was carried out on an EASY-Spray column (50 cm × 75 μm i.d., PepMap C18, 2 μm particles, 100 Å pore size, Thermo Fisher Scientific). Mobile phase A consisted of water and 0.1% v/v formic acid. Mobile phase B consisted of 80% v/v acetonitrile and 0.1% v/v formic acid. Peptides were loaded at a flow rate of 0.3 μl/min and eluted at 0.2 μl/min using a linear gradient going from 2% mobile phase B to 40% mobile phase B over 109 followed by a linear increase from 40% to 95% mobile phase B in 11 min. The eluted peptides were directly introduced into the mass spectrometer. MS data were acquired in the data-dependent mode with a 3 s acquisition cycle. Precursor spectra were recorded in the Orbitrap with a resolution of 120,000. The ions with a precursor charge state between 3+ and 8+ were isolated with a window size of 1.6 m/z and fragmented using high-energy collision dissociation (HCD) with a collision energy of 30. The fragmentation spectra were recorded in the Orbitrap with a resolution of 15,000. Dynamic exclusion was enabled with single repeat count and 60 s exclusion duration. The mass spectrometric raw files were processed into peak lists using ProteoWizard (version 3.0.6618) (Kessner et al., 2008), and cross-linked peptides were matched to spectra using Xi software (version 1.6.745) (Mendes et al., 2018) with in-search assignment of monoisotopic peaks (Lenz et al., 2018). Search parameters were MS accuracy, 3 ppm; MS/MS accuracy, 10ppm; enzyme, trypsin; cross-linker, EDC; max missed cleavages, 4; missing mono-isotopic peaks, 2; fixed modification, carbamidomethylation on cysteine; variable modifications, oxidation on methionine; fragments, b and y ions with loss of H2O, NH3 and CH3SOH.

### Cell culture and transfections

U2OS cells containing 200 copies of an array of 256 tandem repeats of the 17 bp LacO sequence on chromosome 1 (gift from B.E. Black, University of Pennsylvania, Philadelphia; (Janicki et al., 2004)) were grown in DMEM supplemented with 10% FBS and 1% penicillin– streptomycin at 37°C in a 5% CO_2_ incubator. Cells were seeded in 10 cm dishes, a day prior to transfection at a density of 2.5×10^6^ cells per well. Transfections were performed with Lipofectamine 3000 (Life Technologies) according to the manufacturer’s instructions, using 15 μg of plasmid DNA and Opti-MEM I reduced serum medium (Life Technologies). Next day, cells were washed once with 1xDPBS, trypsinised, counted and re-plated on poly-lysine coated coverslips in 6 well plates at a density of 10^6^ cells per well. Downstream experiments were performed three days post-transfection.

### Immunofluorescence

Cells were washed once in PBS and then fixed with 3.7% formaldehyde in 0.1% Triton X-100 in 1xPBS (PBST) for 8 min at RT. Following fixation, the slides were washed once in PBST and then blocked in Image-iT® FX signal enhancer in a humidified chamber at RT for at least 30 min. All antibodies were incubated in a 1:1 mix of PBST and 10% normal goat serum (Life Technologies) overnight at 4°C in a humidified chamber and were used in 1:100 dilution unless otherwise stated: myc (Abcam-ab9106), V5 (Invitrogen-R96025), HA (clone 3F10; E. Kremmer, 1:20). Secondary antibodies coupled to Alexa Fluor 555 and 647 (Invitrogen) were used at 1:100 dilutions. Counterstaining of DNA was performed with DAPI (5 μg/ml) and coverslips were mounted on the slides with 30 μl of SlowFade® Gold antifade reagent (Life technologies)

### Microscopy and image analysis

All immunofluorescence images were taken as 50 z-stacks of 0.2 μm increments, using a 100x oil immersion objective on a Deltavision RT Elite Microscope and a CoolSNAP HQ Monochrome camera. All images were deconvolved using the aggressive deconvolution mode in SoftWorx Explorer Suite (Applied Precision) and are shown as quick projections of maximum intensity.

The mean fluorescence intensity of the protein of interest was measured at the LacO spot, and then the mean fluorescence intensity in the nucleus (background) was subtracted from this value. 25-50 cells were analysed per biological replicate and a minimum of three independent biological replicates were quantified per experiment.

### Analytical ultracentrifugation (AUC)

Sedimentation velocity (SV) and sedimentation equilibrium (SE) experiments were performed using a Beckman Coulter XL-I analytical ultracentrifuge equipped with an An-50 Ti eight-hole rotor. Depending on their concentration, samples were loaded into 12 (low concentration) or 3 mm (high concentration) pathlength charcoal-filled epon double-sector centrepieces, sandwiched between two sapphire windows. For SV, samples were equilibrated at 4°C in vacuum for 6 h before running at 49k rpm. For SE, data were recorded at 26 k rpm. The laser delay, brightness and contrast were pre-adjusted at 3k rpm to acquire the best quality interference fringes. Data were collected using Rayleigh interference and absorbance optics recording radial intensity or absorbance at 280 nm. For SV, data were recorded between radial positions of 5.65 and 7.25 cm, with a radial resolution of 0.005 cm and a time interval of 7 minutes, and analysed with the program SEDFIT (Schuck, 2000) using a continuous c(s) model. For SE data were recorded between radial positions of 6.00 and 7.25 cm, with a radial resolution of 0.001 cm and a time interval of 3 h (until successive scans overlaid satisfactorily), and analysed with the program SEDPHAT (Vistica et al., 2004) using species analysis. The partial specific volume, buffer density and viscosity were calculated using SEDNTERP (Hayes et al., 2012). Sedimentation coefficients were computed from atomic coordinate models using SOMO (Brookes and Rocco, 2018).

## DATA AND SOFTWARE AVAILABILITY

The accession numbers for the coordinates and structure factors reported in this paper are PDB: xxxx-CAL1_1-160_–CENP-A/H4 form I, xxxx-CAL1_1-160_–CENP-A/H4 form II, xxxx-CENP-C_1264-1411_ and xxxx-CENP-C_1264-1411_-CAL_1841-979_. Crosslink date is deposited in y.

