## Supplemental Figures for "Structural Basis for CAL1-Mediated Centromere Maintenance"

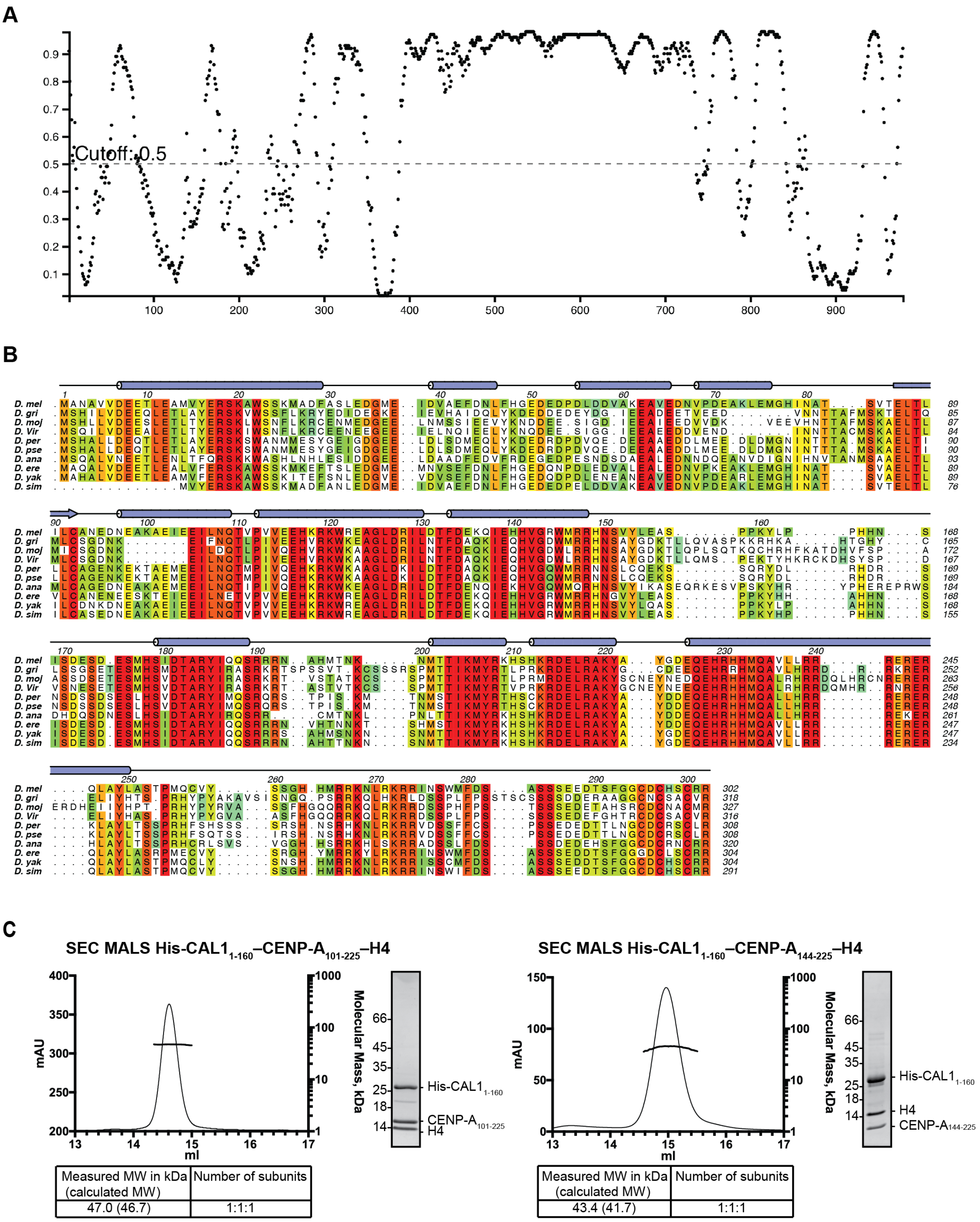


**Figure S1. Related to Figure 1. CAL1 is predicted to be predominantly unstructured**

A - Disordered regions of CAL1 as predicted by Disopred (http://bioinf.cs.ucl.ac.uk/psipred).

B - Secondary structure composition and multiple sequence alignment of CAL1 N-terminus as performed using Psipred (http://bioinf.cs.ucl.ac.uk/psipred) and with MUSCLE (Madeira et al., 2019). Numbers correspond to *D. mel*. *D. melanogaster* (*D. mel*), *D. grimshawi* (*D. gri*), *D.*

*mojavensis* (*D. moj*), *D. virilis* (*D. viri*), *D. persimilis* (*D. per*), *D. pseudoobscura pseudoobscura* (*D. pse*), *D. erecta* (*D. ere*), *D. yakuba* (*D. yak*) and *D. simulans* (*D. sim*).

C - SEC-MALS analysis of His-CAL1$_{1\text{-}160}$–CENP-A$_{101\text{-}225}$–H4 and His-CAL1$_{1\text{-}160}$–CENP-A$_{144\text{-}225}$–H4. Absorption at 280 nm (mAU, left y-axis) and molecular mass (kDa, right y-axis) are plotted against elution volume (ml, x-axis). Measured molecular weight (MW) and the calculated subunit stoichiometry based on the predicted MW of different subunit compositions. Samples run on Superdex 200 increase 10/300 in 50 mM HEPES pH8.0, 2 M NaCl and 1 mM TCEP.

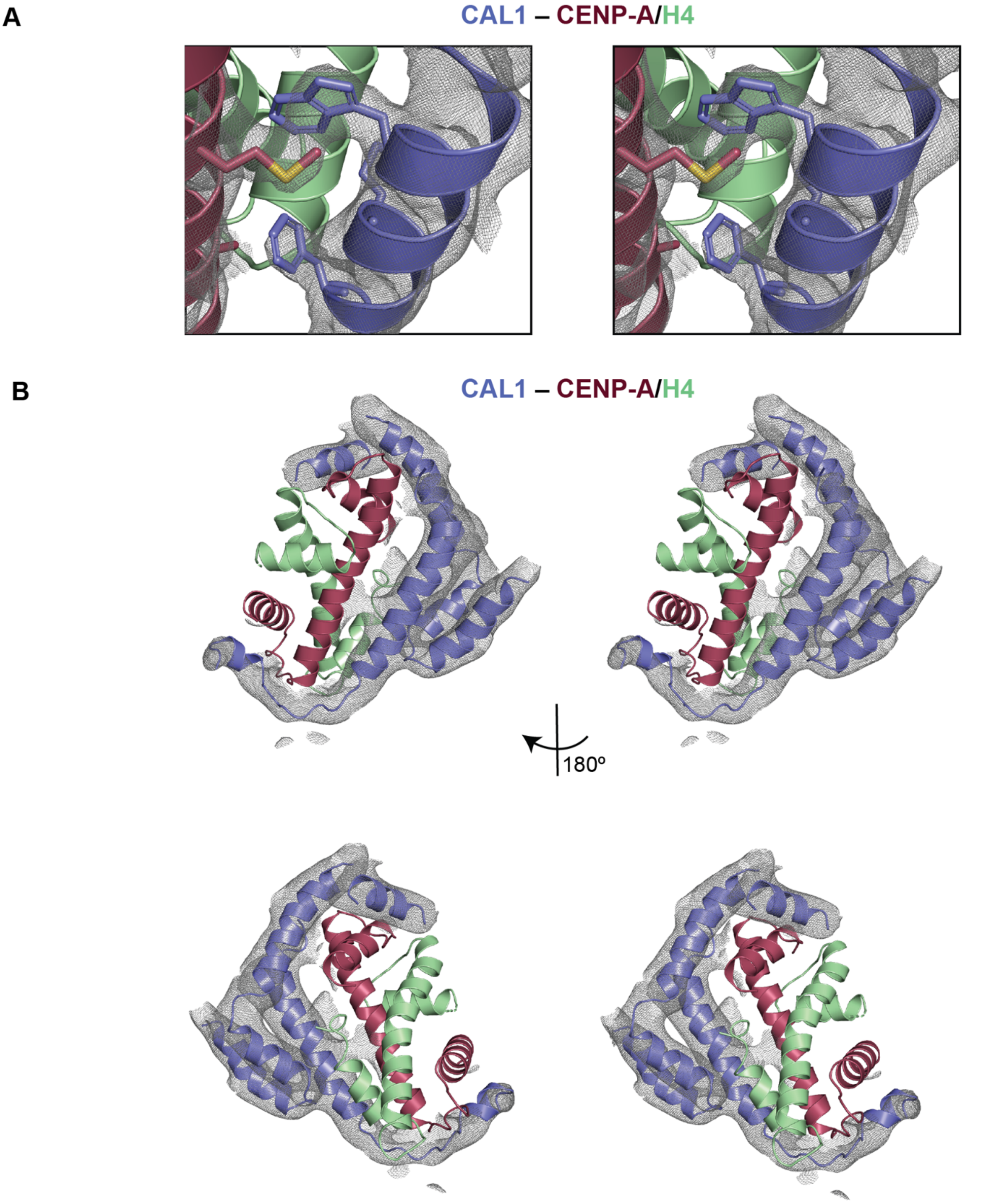


**Figure S2. Related to Figure 1. Stereo images of the electron density maps and corresponding final models of CAL1–CENP-A/H4 from crystal form I and II**

A - 2Fo-Fc electron density map contoured at 1 sigma for the vicinity of CAL1 residues W22 and F29 for crystal form I. CENP-A in maroon and H4 in green.

B - 2Fo-Fc electron density map contoured at 1 sigma for CAL1 bound to CENP-A/H4 for crystal form II. CAL1 is shown in blue, CENP-A in maroon and H4 in green.

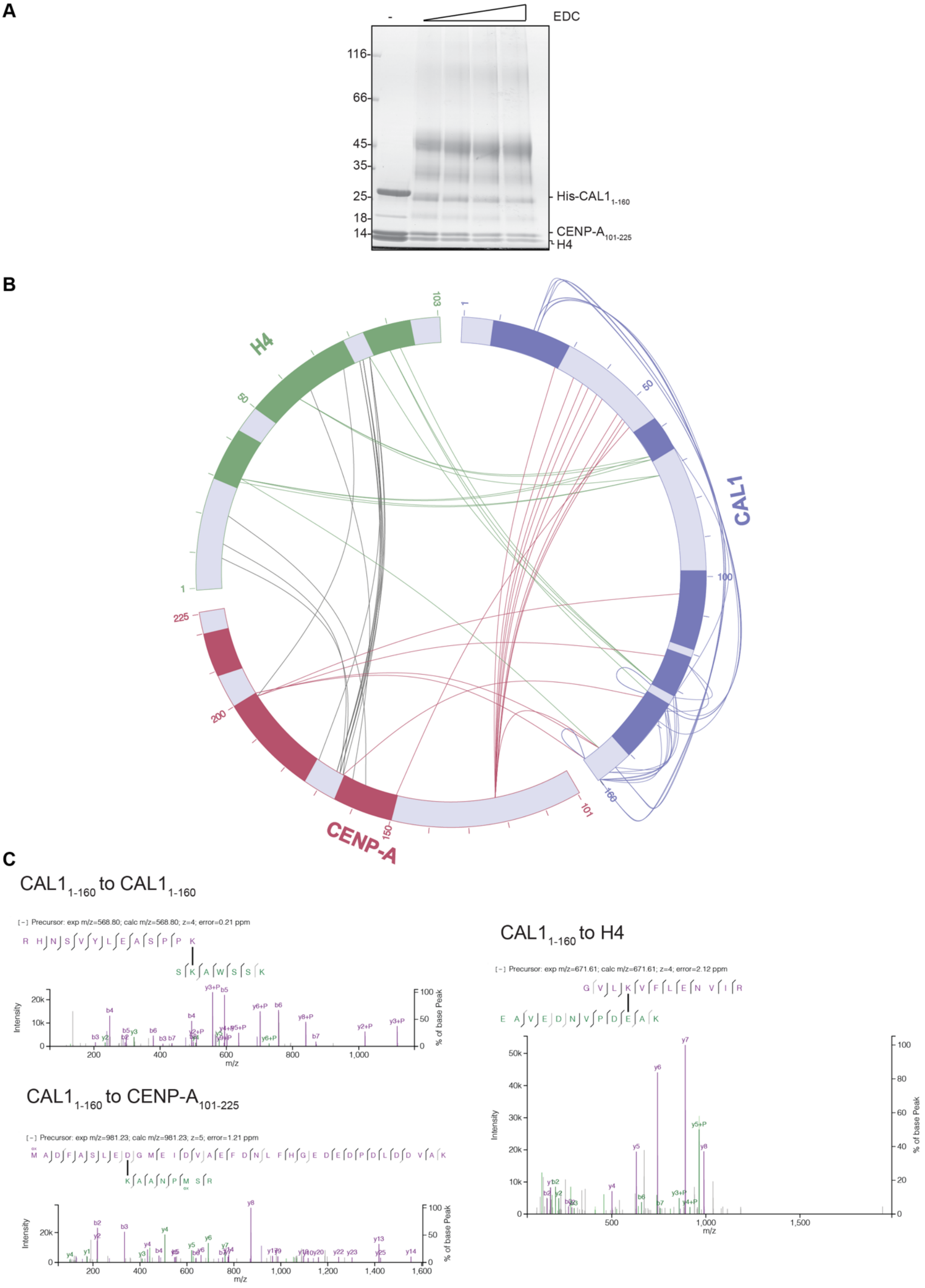

A
EDC
116
66
45
35
25
18
14
His-CAL1 1-160
CENP-A 101-225
H4
B
H4
CAL1
CENP-A
103
1
50
100
160
225
200
150
101
C
CAL1 1-160 to CAL1 1-160
[−] Precursor: exp m/z=568.80; calc m/z=568.80; z=4; error=0.21 ppm
R H N S V Y L E A S P P K
S K A W S S K
Intensity
% of base Peak
m/z
CAL1 1-160 to CENP-A 101-225
[−] Precursor: exp m/z=981.23; calc m/z=981.23; z=5; error=1.21 ppm
M A D F A S L E D G M E I D V A E F D N L F H G E D E D P D L D D V A K
K A A N P M S R
Intensity
% of base Peak
m/z
CAL1 1-160 to H4
[−] Precursor: exp m/z=671.61; calc m/z=671.61; z=4; error=2.12 ppm
G V L K V F L E N V I R
E A V E D N V P D E A K
Intensity
% of base Peak
m/z

**Figure S3. Related to Figure 1. Intra and Intermolecular contacts identified between CAL1 and CENP-A/H4 using Cross-Linking combined Mass Spectrometry (CLMS)**

A - SDS-PAGE analysis of His-CAL1$_{1\text{-}160}$-CENP-A$_{101\text{-}225}$-H4 crosslinked with increasing amounts of EDC cross linker.

B - Linkage map showing the sequence position and crosslinked residues pairs between His-CAL1$_{1\text{-}160}$–CENP-A$_{101\text{-}225}$–H4. EDC crosslinked samples were resolved with SDS-PAGE then analysed by MS. CAL1 is shown in blue, CENP-A in maroon and H4 in green.

C - High resolution representative fragmentation spectra displayed using XiSpec (Ref: PMID: 29741719) for crosslinked peptides seen between CAL1$_{1\text{-}160}$ and CAL1$_{1\text{-}160}$, CAL1$_{1\text{-}160}$ and CENP-A$_{101\text{-}225}$ and CAL1$_{1\text{-}160}$ and H4

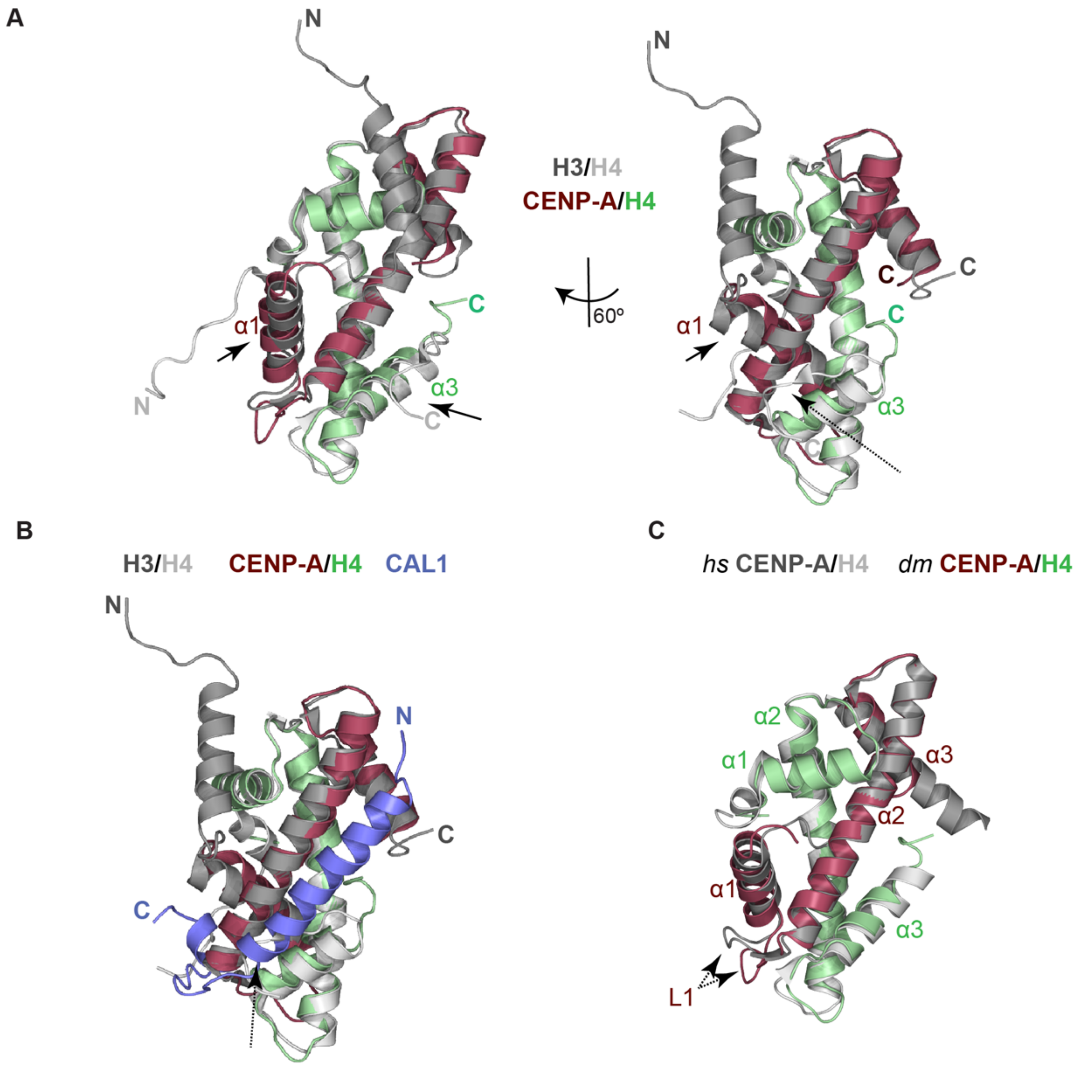


**Figure S4. Related to Figure 1. Mode of H4 recognition by CENP-A is similar to H3.**

A - Structural superposition of CENP-A/H4 with H3/H4. CENP-A is shown in maroon, H4 in green and H3/H4 in silver. Arrows indicate conformational changes.

B - Structural superposition of CAL1–CENP-A/H4 with H3/H4. CAL1 is shown in blue, CENP-A in maroon, H4 in green and H3/H4 in silver. Arrows indicate conformational changes; dotted arrow highlights conformational changes in the loop regions.

C - Structural superposition of *hs* CENP-A/H4 with *dm* CENP-A/H4. *dm* CENP-A in maroon, *dm* H4 in green and *hs* CENP-A/H4 in silver. Dotted arrow highlights conformational changes in the loop regions.

A

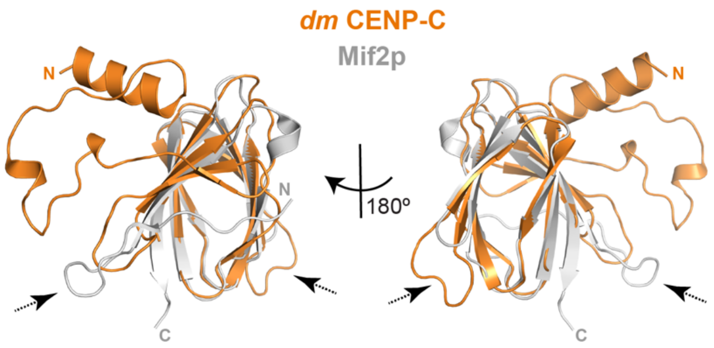


B

| Species | Mass (g/mol) | | $s^{0}_{20,W}$ (S) | |
|---|---|---|---|---|
| | Calculated | Experimental | Computed | Experimental |
| $CAL1_{841-979}$ | 15,391 | 19,111 | 1.48 | 1.46 |
| $(CAL1_{841-979})_2$ | 30,782 | | 2.02 | |
| $CENP\text{-}C_{1264-1411}$ | 17,187 | 33,867 | 1.77 | 2.74 |
| $(CENP\text{-}C_{1264-1411})_2$ | 34,374 | | 2.96 | |
| $(CENP\text{-}C_{1264-1411})_2 + CAL1_{841-979}$ | 49,765 | 48,889 | 3.47 | 3.50 |
| $(CENP\text{-}C_{1264-1411})_2 + (CAL1_{841-979})_2$ | 65,156 | | 3.89 | |

**Figure S5. Related to Figure 5 and Figure 7. Dimeric CENP-C binds one copy of CAL1.**

A - Structural superposition of cupin domains of *dm* CENP-C and Mif2p. CENP-C is shown orange and Mif2p in silver. Dotted arrow highlights conformational changes in the loop regions.

B - Calculated and experimentally determined molecular masses and sedimentation coefficients for plausible solution states of $CAL1_{841-979}$, $CENP\text{-}C_{1264-1411}$ and complexes thereof.

**Table S1**

| | $CAL1_{1-160}$–CENP-A/H4 Form I | $CAL1_{1-160}$–CENP-A/H4 Form II | CENP-$C_{1264-1411}$ | $CAL1_{841-979}$–CENP-$C_{1264-1411}$ |
|---|---|---|---|---|
| **Data Collection** | | | | |
| Space group | R 3 2 :H | P 63 2 2 | P 41 21 2 | P 21 21 21 |
| **Cell dimensions** | | | | |
| *a*, *b*, *c*, (Å) | 178.25, 178.25, 133.28 | 199.70, 199.70, 76.75 | 57.20, 57.20, 92.96 | 86.27, 86.44, 88.46 |
| α, β, γ (°) | 90, 90, 120 | 90, 90, 120 | 90, 90, 90 | 90, 90, 90 |
| Wavelength | 0.97625 | 0.91587 | 0.97623 | 0.97625 |
| Resolution (Å) | 100.9 - 3.47 (3.59 - 3.47) | 70.16 - 4.36 (5.19 - 4.37) | 28.6 - 1.82 (1.88 - 1.82) | 29.08 - 2.27 (2.35 - 2.27) |
| $R_{merge}$ | 0.147 (1.96) | 0.129 (0.95) | 0.0548 (0.598) | 0.076 (0.663) |
| $R_{pim}$ | 0.058 (0.759) | 0.046 (0.526) | 0.013 (0.177) | 0.022 (0.202) |
| $I/\sigma I$ | 10.46 (1.15) | 9.61 (2.4) | 34.13 (3.55) | 21.16 (3.40) |
| Completeness (%) | 99.93 (100.00) | 38 (8) | 99.73 (97.87) | 98.41 (94.61) |
| Completeness (ellipsoidal) (%) | | 89 (81) | | |
| Redundancy | 7.6(7.7) | 8.9 (8.8) | 18.0 (11.9) | 13.1 (11.3) |
| **Refinement** | | | | |
| No. reflections | 80,934 (8111) | 20,689 (307) | 14,426 (1381) | 404,275 (33273) |
| $R_{work}$ (%) /$R_{free}$ (%) | 25.5 / 28.4 | 30.8 / 32.7 | 19.5 / 23.2 | 23.0 / 26.9 |
| **No. atoms** | | | | |
| Protein | 2781 | 1746 | 1085 | 4235 |
| ***B*-factors** | | | | |
| Protein | 138.3 | 232 | 42.9 | 72.3 |
| **R.m.s deviations** | | | | |
| Bond length (Å) | 0.010 | 0.004 | 0.007 | 0.005 |
| Bond angles (°) | 1.1 | 1.20 | 0.94 | 1.10 |
| **Ramachandran values** | | | | |
| Favored (%) | 90.6 | 89.32 | 97.79 | 99.23 |
| Dissallowed (%) | 2.9 | 1.7 | 0.00 | 0.0 |

Data collection and refinement statistics
Statistics for the highest-resolution shell are shown in parentheses.

**Table S2**

Primers used during this study

| Construct | Forward | Reverse |
|---|---|---|
| pEC-K-3C-His CAL1$_{1\text{-}160}$ | 5'-ccaggggcccgactcgatggctaacgcggttg-3' | 5'-cagaccgccaccgactgcttattttggcgggctcgc-3' |
| Codon Optimised CAL1$_{1\text{-}160\ W22A/F29A}$ | 5'-acgaacgttccaaagctgcgtcctctaagatggcgg-3'<br>5'-ggtcctctaagatggcggatgctgcatccctggaagat-3' | 5'-ccgccatcttagaggacgcagctttggaacgttcgt-3'<br>5'-atcttccagggatgcagcatccgccatcttagaggacc-3' |
| Codon Optimised CAL1$_{1\text{-}160\ W22R/F29R}$ | 5'-ggtatacgaacgttccaaagctaggtcctctaagatg-3'<br>5'-ggtcctctaagatggcggatcgtgcatccctggaagat-3' | 5'-catcttagaggacctagctttggaacgttcgtatacc-3'<br>5'-atcttccagggatgcacgatccgccatcttagaggacc-3' |
| Codon Optimised CAL1$_{1\text{-}160\ F43R}$ | 5'-gcatggaaattgacgtcgcagaacgcgataacctgttccacgg-3' | 5'-ccgtggaacaggttatcgcgttctgcgacgtcaatttccatgc-3' |
| pEC-K-3C-His CAL1$_{1\text{-}50}$ | 5'- ccaggggcccgactcgatggctaacgcggttg-3' | 5'-cagaccgccaccgactgcttattcaccgtggaacagg-3' |
| 14S CAL1$_{841\text{-}979}$ | 5'-tacttccaatccaatgcaatgggtggtgacccggattg-3' | 5'-ttatccacttccaatgttatta cttatcaccggagttg-3' |
| Codon Optimised CAL1$_{841\text{-}979\ I900R/K907A/Y908A}$ | 5'-acagggcctgggcaaaatcaggggcgaacgttg-3'<br>5'aatcatcggcgaacgttgggcgcgtgcggccctgaaataccacattgg tagccgttcctt-3' | 5'-caacgttcgcccctgattttgcccaggccctgt-3'<br>5'aaggaacggctaccaatgtggtatttcagggccgcacgcgcccaa cgttcgccgatgatt-3 |
| Non-Codon Optimised CAL1 $_{W22A/F29A}$ | 5- gagcgctctaaggccgcgtccagtaaaatggcg-3'<br>5'-ctggtccagtaaaatggcggatgctgccagcctggaa-3' | 5'-cgccattttactggacgcggccttagagcgctc-3'<br>5'-ttccaggctggcagcatccgccattttactggaccag-3' |
| Non-Codon Optimised CAL1 $_{W22R/F29R}$ | 5'-gagcgctctaaggccaggtccagtaaaatgg-3'<br>5'-ctggtccagtaaaatggcggatcgtgccagcctggaa-3' | 5'-ccattttactggacctggccttagagcgctc-3'<br>5'-ttccaggctggcacgatccgccattttactggaccag-3' |
| Non-Codon Optimised CAL1 $_{F43R}$ | 5'-gaaatagatgtggccgagcgcgacaacttgttccacgg-3' | 5'-ccgtggaacaagttgtcgcgctcggccacatctatttc-3' |
| Non-Codon Optimised CAL1 $_{I900R/K907A/Y908A}$ | 5'-cgagcagggactcggaaagattaggggagaacgttggg-3'<br>5'-ggagaacgttgggcccgcgcggccctgaagtaccacatcgg-3" | 5'-cccaacgttctcccctaatctttccgagtccctgctcg-3'<br>5'ccgatgtggtacttcagggccgcgcgggcccaacgttctcc-3' |
| CENP-A$_{101\text{-}225\ S154Q}$ | 5'-ccaagctgccgttccagcgtctagtgcgcg-3' | 5'-cgcgcactagacgctggaacggcagcttgg-3' |
| CENP-A$_{101\text{-}225\ M186A}$ | 5'-caggagtcgtgcgaggcgtacttgacgcagcg -3' | 5'-cgctgcgtcaagtacgcctcgcacgactcctg -3' |
| CENP-A$_{101\text{-}225\ Q190G}$ | 5'-gagatgtacttgacggggcggctcgccgactc-3' | 5'-gagtcggcgagccgccccgtcaagtacatctc-3' |
| CENP-A$_{101\text{-}225\ M186A\ Q190G}$ | 5'-caggagtcgtgcgaggcgtacttgacggggcg-3' | 5'-cgccccgtcaagtacgcctcgcacgactcctg-3' |
| pEC-K-3C-His CENP-C$_{1264\text{-}1411}$ | 5'-ccaggggcccgactcgatgggcccggtagtgttc-3' | 5'-cagaccgccaccgactgcttaagaacggatacac-3' |
